# IMIX: A multivariate mixture model approach to integrative analysis of multiple types of omics data

**DOI:** 10.1101/2020.06.23.167312

**Authors:** Ziqiao Wang, Peng Wei

## Abstract

**Motivation:** Integrative genomic analysis is a powerful tool to study the biological mechanisms underlying a complex disease or trait across multiplatform high-dimensional data, such as DNA methylation, copy number variation (CNV), and gene expression. It is common to perform large-scale genome-wide association analysis of an outcome for each data type separately and combine the results *ad hoc*, leading to loss of statistical power and uncontrolled overall false discovery rate (FDR).

**Results:** We propose a multivariate mixture model framework (IMIX) that integrates multiple types of genomic data and allows examining and relaxing the commonly adopted conditional independence assumption. We investigate across-data-type FDR control in IMIX, and show the gain in lower misclassification rates at controlled overall FDR compared with established individual data type analysis strategies, such as Benjamini-Hochberg FDR control, the q-value, and the local FDR control by extensive simulations. IMIX features statistically-principled model selection, FDR control, and computational efficiency. Applications to the Cancer Genome Atlas (TCGA) data provide novel multi-omic insights into the luminal/basal subtyping of bladder cancer and the prognosis of pancreatic cancer.

**Availability and implementation:** We have implemented our method in R package “IMIX” with instructions and examples available at https://github.com/ziqiaow/IMIX.

## 1 Introduction

With the development of high-throughput technology, integrative genomic analysis has become a powerful tool in biomedical research to study the biological mechanisms underlying a complex disease or trait. The mechanisms behind a certain disease outcome, such as the prognosis or the molecular subtypes of cancer, involve alterations in multiple pathways and biological processes including copy number variations (CNV), epigenetic changes, transcriptomic changes. It is a challenge to integrate all this information together to analyze the disease outcome. A common strategy is to assess the associations between genes and an outcome separately for each data type using Bonferroni correction or the Benjamini-Hochberg false discovery rate (BH-FDR) procedure (Benjamini and Hochberg, 1995) to adjust for multiple hypothesis testing. However, study on The Cancer Genome Atlas (TCGA) has found that omics data, such as gene expression, DNA methylation, and CNV, have several different triangular dependence structures (Sun et al., 2018). Furthermore, it remains unclear whether the correlation structures between data types vary according to the associations between different genes and the outcome of interest through those data types. Therefore, the heuristic separate analysis strategy loses statistical power by assuming that the data sources are independent of each other. The integration of multiple types of omics data by identifying the unknown dependence structures becomes essential to understand the intricacy of the genomic mechanisms behind complex diseases.

Previous developments in integrative genomics models mainly solve two types of problems: exploration (unsupervised) and prediction (supervised); see Richardson et al. (2016) for a comprehensive review. Unsupervised methods, such as clustering, focus on identifying similarity and uncovering heterogeneity among individuals, accounting for intersource associations between multiple data types (Kirk et al., 2012; Mo et al., 2013; Shen et al., 2009). These methods are exploratory techniques rather than hypothesis-testing tools. Supervised methods, such as prediction or regression analysis, concentrate on outcome prediction and feature selections, such as penalized regression analysis (Tibshirani, 1996; Zou and Hastie, 2005) and nonparametric-based approach (Pineda et al., 2015). While these methods consider the association between genes and the outcome, to our best knowledge, most of them require individual-level data from the same set of samples and/or have no rigorous error-control procedure.

Here we propose a multivariate mixture model approach (IMIX) to integratively analyze omics data using summary statistics that incorporates the correlations and biological coordinations between multiple data sources to boost the statistical power for genomic discovery while controlling the across-data-type FDR. We use the EM algorithm to estimate the model parameters and propose an adaptive FDR control procedure. IMIX also features statistically-principled model selection and does not require the use of a common set of samples across data types, which relaxes the conditions of previously published methods for the integration of multiple omic data.

Through extensive simulation studies, we demonstrated that IMIX yielded better statistical power and overall FDR control compared with individual data type analysis strategies, such as BH-FDR, Bonferroni corrections, the q-value, and the local FDR control. We also observed that IMIX is computationally efficient. Lastly, we applied IMIX to study the molecular subtypes of bladder cancer through DNA methylation, CNV, and gene expression, as well as the prognosis of pancreatic cancer through gene expression, and CNV in the TCGA. Our applications of IMIX to the two TCGA data sets showed that different genomic data types can be correlated in both non-disease-associated and disease-associated genes, refuting the commonly adopted conditional independence assumption in integrative analysis of multiplatform genomic data.

## 2 Methods

In this section, we provide some background regarding IMIX (Section 2.1), and consider how these may be extended to allow us to perform integrative modeling of multiple datasets and model selection (Section 2.2). In section 2.3, we discuss model selection regarding the number of mixture components and the best model among proposed variants of the multivariate mixture model. We discuss local FDR (LFDR) and the adaptive procedure for across-data-type FDR control in Section 2.4.

### 2.1 Preliminaries and Notations for IMIX

Consider the association testing problem between gene *i, i* = 1, 2, …, *N* and the outcome through data type *h* = 1, 2, …, *H*. For example, one wants to identify which gene is associated with a binary outcome, basal or luminal molecular subtype of bladder cancer, and assess the associations through *H* = 3 genomic data types: DNA methylation, gene expression, and CNV. It is a common practice to assume *a priori* that each genomic data type is independent and to perform statistical analysis for each dataset separately. Then, one can apply state-of-the-art methods such as the BH-FDR or the q-value for each data type to adjust for multiple hypothesis testing. Here, we group the genes into *K* = 2^*H*^ latent states according to their associations with the *h* data types, and we introduce a vector of binary variables to denote each latent state *k* of gene *i*: ***z***_*ik*_ = (*z*_*ik*1_, *z*_*ik*2_, …, *z*_*ikh*_). Without loss of generality, we assume *H* = 3. When *k* = 1, 2, …, 8, the potential latent states/classes of gene *i* are: ***z***_*i*1_ = (0, 0, 0), ***z***_*i*2_ = (1, 0, 0), ***z***_*i*3_ = (0, 1, 0), ***z***_*i*4_ = (0, 0, 1), ***z***_*i*5_ = (1, 1, 0), ***z***_*i*6_ = (1, 0, 1), ***z***_*i*7_ = (0, 1, 1), and ***z***_*i*8_ = (1, 1, 1). If *z*_*ikh*_ = 1, gene *i* is associated with the outcome through data type *h* in class *k*; if *z*_*ikh*_ = 0, gene *i* is not associated with the outcome through data type *h* in class *k*. Dependeing on the latent state of gene *i*, i.e., whether it belongs to latent state *k* or not, we have *T*_*ik*_ = 1 or *T*_*ik*_ = 0, repsectively.

We assume that the data can be summarized as *x*_*ih*_ for each gene *i, i* = 1, …, *N* and data type *h* = 1, 2, 3. Here, *x*_*ih*_ is a Z-score that is transformed from the p-value of the association test for each data type (McLachlan et al., 2006; Wei and Pan, 2008). The transformation is given by *x*_*ih*_ = Φ^−1^(1 − *p*_*ih*_), where Φ is the cumulative distribution function of the standard normal distribution *N* (0, 1), and *p*_*ih*_ is the p-value for gene *i* and data type *h*.

We assume that *X*_*i*_ = (*x*_*i*1_, *x*_*i*2_, *x*_*i*3_)^*T*^ comes from a mixture distribution with *K* = 8 mixture components:

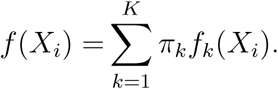

Here, we assume the Z scores of genes in class *k* follow an *H*-dimensional multivariate distribution *f*_*k*_, and the mixing proportions are *π*_*k*_, *k* = 1, *…*, 8. To assess how likely gene *i* belongs to the latent state *k*, we estimate the posterior probability of the latent label *T*_*ik*_:

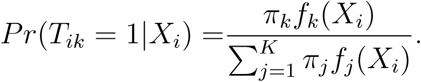

### 2.2 Parametric Multivariate Mixture Model

Based on the principles of molecular biology and previous publications (Sun et al., 2018; Wei and Pan, 2012), when a gene is associated with an outcome through, for example, DNA methylation and gene expression, the two events can be correlated. In particular, Wei and Pan (2012) developed an integrative genomic method to improve the power of a two-component mixture model by considering the possible correlations between three data types. Here we extend this idea and formulate a multivariate mixture model of *K* = 2^*H*^ components. We assume the *k*-th component distribution *f*_*k*_ to be multivariate normal. Normal mixture model can be used and is widely used to approximate many mixture distributions in real data applications (Sun and Cai, 2007). The marginal mixture density *f* (***X***_***i***_) can then be written as

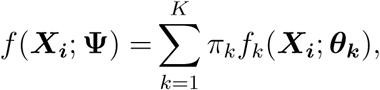

where

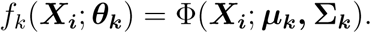

Here, the vector Ψ = (*π*_1_, *π*_2_, *…, π*_*K*−1_, ***ξ***^*T*^)^*T*^ contains all the unknown parameters in the mixture model. ***ξ*** is the vector containing all the elements of the component means, ***µ***_**1**_, *…*, ***µ***_***K***_, and the elements of the covariance matrices, **Σ**_**1**_, *…*, **Σ**_***K***_, known *a priori* to be distinct. We use the expectation-maximization (EM) algorithm (Dempster et al., 1977) to estimate **Ψ**. We call this generic multivariate Gaussian mixture model as **IMIX-Cor**. In the following sections we will describe three variants based on this model.

#### 2.2.1 IMIX-Cor-Restrict: Correlated Mixture Model with Restrictions on Mean

To tackle the possible unidentifiability problem of **Ψ** due to the interchanging of component labels, we impose the following constraints on ***µ***_*k*_ :

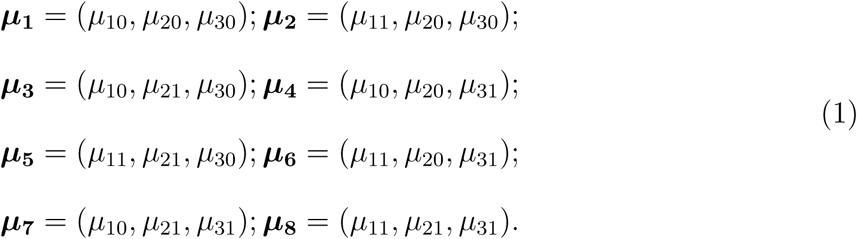

These constraints correspond to the biological rational that for each data type, we assume that the means of the test statistics from the null and non-null groups are the same across the *K* classes. For example, *µ*_10_, the mean of the null group in data type 1, is the same in ***µ***_**1**_ and ***µ***_**3**_. The multivariate Gaussian mixture model with resctrictions on the mean is denoted as **IMIX-Cor-Restrict**. We estimate the parameters using the EM algorithm (Supplementary Materials, Section 1).

#### 2.2.2 IMIX-Ind: Independent Mixture Model with Restrictions on Mean and Variance

If we assume there is no correlation between any two data types, then the covariance matrix in IMIX-Cor-Restrict, Σ_*k*_, becomes diagonal. The model is reduced to

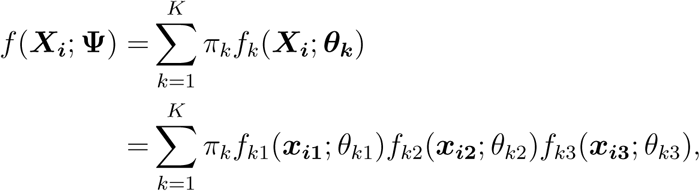

where *f*_*kh*_(***x***_***ih***_; *θ*_*kh*_) = Φ(***x***_***ih***_; *µ*_*kh*_, *σ*_*kh*_) is a normal probability density function with mean *µ*_*kh*_ and variance 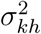, *k* = 1, *…*, 8; *h* = 1, 2, 3. Besides the constraints on the mean in (1), we impose the same constraints on the variance 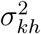 on the basis of the null and non-null genes for each data type. We call this model **IMIX-Ind**.

#### 2.2.3 IMIX-Cor-Twostep: Correlated Mixture Model with Fixed Mean

To reduce the model complexity, i.e., to create a more parsimonious model, and to ease the computation time, we propose a third modification based on the previous models, the correlated mixture model with fixed mean. This model is similar to the previous model with constraints on the means; however, with the replacement of the means estimated from the independent model IMIX-Ind, we ease the complexity of estimating the mean and the covariance matrices at the same time in the EM algorithm. Based on our simulation study to be detailed later on, IMIX-Ind performed well in estimating the means; thus, to facilitate the correlation estimation between data types, we introduce the correlated mixture model with fixed mean, where we only estimate the covariance matrix for each component with a pre-specified mean vector. This provision would ease the time from estimating both the mean and the covariance together; we will show later in the simulation study that this model achieves the best computational efficiency among IMIX models that consider the correlation structures. Here, the model estimation follows the same EM procedure as before. We call this model **IMIX-Cor-Twostep**.

### 2.3 Model Selection

In real data applications, one or a few classes out of *K* may be absent, for example, there may be no gene in class 8 in the TCGA bladder cancer example, where no gene was associated with the outcome through all three molecular mechanisms. Therefore, elimination of unnecessary components is crucial, which leads to more precise parameter estimation and well-calibrated FDR control. This is closely related to the question of how many components *K* to include in the mixture distribution to prevent overfitting. As pointed out by previous works (Leroux, 1992; McLachlan and Peel, 2004), the penalized log likelihood functions including Akaike Information Criterion (AIC) and Bayesian Information Criterion (BIC) are adequate for the number of components selection under a finite mixture distribution; in particular, under mild conditions, AIC and BIC do not underestimate the true number of components asymptotically. Specifically, AIC and BIC select the model that, respectively, minimizes

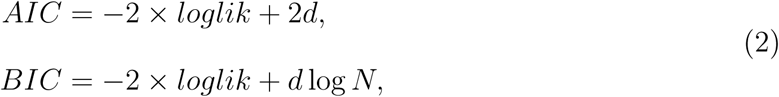

where *d* is the number of unknown parameters (i.e., degrees of freedom), *N* is the number of genes, and loglik is the maximized full log likelihood.

Along with model selection for the number of components *K*, we also select the best model for a fixed *K* among the different methods introduced in Section 2.2 regarding mean and covariance structures. We introduce the IMIX framework, where the data is fitted for all four IMIX methods (IMIX-Ind, IMIX-Cor, IMIX-Cor-Twostep, and IMIX-Cor-Restrict), then AIC or BIC is utilized to select the best model among a number of candidate models, called IMIX-AIC or IMIX-BIC.

### 2.4 Local FDR Control and Adaptive Procedure for across-datatype FDR Control

For the trivariate mixture model of *H* = 3 data types, there are *K* = 8 latent classes, and we classify each gene *i* to component 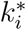, based on 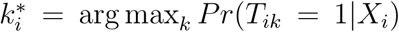. The hypothesis to be tested is

*H*_0,*i*_ : Gene *i* does not belong to component 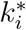;

*H*_1,*i*_ : Gene *i* belongs to component 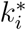.

The estimated posterior probability that gene *i* belongs to 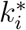 is defined as 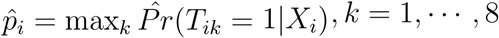. The LFDR for gene *i* is defined as 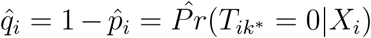. If 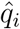 is smaller than a threshold *t*, we reject *H*_0,*i*_. This enables us to control LFDR according to the correspondence between the posterior probability and the threshold *t* (Efron et al., 2007).

In addition to controlling LFDR, Sun and Cai (2007) proposed an adaptive procedure to control the global FDR for mixture models. We propose to extend their method to the IMIX framework. For each component *k*, we construct the following hypothesis

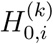 : Gene *i* does not belong to component *k*;

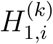 : Gene *i* belongs to component *k*.

Our method controls the across-data-type FDR for component *k* and it is defined as FDR_*k*_=*E*(*N*_10,*k*_*/R*_*k*_|*R*_*k*_ *>* 0)*Pr*(*R*_*k*_ *>* 0), *k* = 1, …, *K*. Here *N*_10,*k*_ is the number of false discoveries in component *k* and *R*_*k*_ is the total number of hypotheses claimed significant in component *k*. When no hypothesis is claimed significant, FDR_*k*_ is 0. The estimated posterior probability that gene *i* belongs to component *k* is defined as 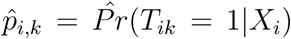, *k* = 1, …, 8. The local FDR (LFDR) for gene *i* is defined as 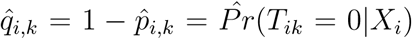. The adaptive step-up procedure is described below:

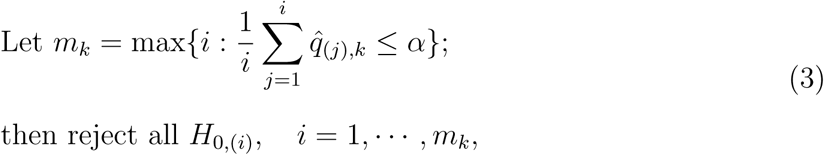

where 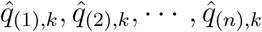 are the ranked values of the LFDR in component *k*. The adaptive procedure controls the marginal FDR (mFDR) for each component at level *α* asymptotically. Here mFDR_*k*_ is defined as *E*(*N*_10,*k*_)*/E*(*R*_*k*_).The estimated mFDR becomes 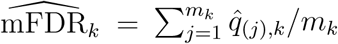. Genovese and Wasserman (2002) showed that under weak conditions, there exists an asymptotic relationship between mFDR and the across-data-type FDR of one component, in which mFDR_*k*_ = FDR_*k*_ + *O*(*N* ^−1*/*2^), where *N* is the number of hypotheses in component *k* and it is the same across all components. This adaptive procedure can be further used for a combination of components. For example, if we are interested in all the genes that are associated with the outcome through DNA methylation in a three-data type integration problem of DNA methylation (M), gene expression (E), and CNV, the procedure can be applied to the combination of component 2, 5, 6, and 8, i.e., (M+,E-,CNV-), (M+,E+,CNV-), (M+,E-,CNV+), and (M+,E+,CNV+).

## 3 Results

### 3.1 Simulation Studies

We performed two sets of simulation studies. Simulation study 1 assessed the performance of IMIX in terms of across-data-type FDR control, misclassification rate, and model calibration; simulation study 2 assessed the information criteria we proposed for model selection. We consider the following multivariate normal mixture model for three data types:

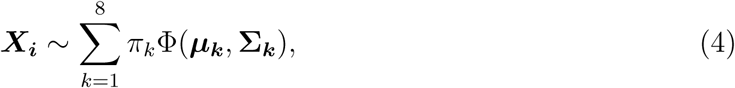

where ***µ***_***k***_ = (*µ*_*k*1_, *µ*_*k*2_, *µ*_*k*3_) with *µ*_*kh*_ corresponds to the mean of data type *h* in component *k*, and **Σ**_***k***_ is a 3 × 3 matrix that contains the variance 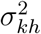 and the covariance *σ*_*khh*′_ between data type *h* and *h*′ in component *k*.

#### 3.1.1 Across-data-type FDR Control and Misclassification Rate

To illustrate the gain in lower misclassification rate while controlling for the across-data-type FDR and LFDR of IMIX compared with the commonly used methods, we generated 1 000 simulated datasets of *N* × *p* = 20 000 × 3 Z-scores *x*_*ih*_ following (4) in six scenarios. Scenario 1 assumed all three data types were independent with **Σ**_***k***_ = diag(1,1,1); the mean under the null was 0 and under the alternative was 3 for data type h; the proportion of each component was balanced as *π*_*k*_ = 0.125. Scenarios 2-5 assumed the Z-scores were correlated under the alternative hypothesis by adding covariances (here they were also the correlations) *σ*_*k*12_ = *σ*_*k*13_ = *σ*_*k*23_ in **Σ**_***k***_; here we only set the covariances to be non-zero for *k* = 5, …, 8, and each scenario corresponded to a covariance of 0.1, 0.3, 0.5, and 0.8, respectively. The rest of the parameters in Scenario 2-5 followed those of Scenario 1. Scenario 6 mimicked the real data in Section 3.2.1, where we analyzed the luminal/basal molecular subtypes through DNA methylation, gene expression, and CNV of the bladder cancer data in TCGA. We set the mean and covariance matrices in (4) equal to the empirical values estimated from Z-scores classified by separate analysis of each data type using BH-FDR method, and an unbalanced proportion equal to the estimated 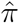 using IMIX-Cor-Twostep (Supplementary Materials, Section 2.1). This simulation scenario thus did not favor either the separate analysis or the IMIX method.

We analyzed the simulated data by applying our proposed methods, including IMIX-Ind, IMIX-Cor-Restrict, IMIX-Cor, IMIX-Cor-Twostep, IMIX-AIC, and IMIX-BIC; to compare the model performance with commonly used separate analysis methods, we also applied BH-FDR (Benjamini and Hochberg, 1995), Bonferroni correction, q-value (Storey, 2002; Storey et al., 2019), and the local FDR procedure (Efron et al., 2007). We set *α* = 0.2 as the nominal error control level across all methods for comparisons. Note that we suggest an *α* value threshold between 0.05 and 0.2 for IMIX to discover interesting non-null genes while controlling the proportion of null genes (false positives) in the significant gene list.

The simulation results are presented in Figure 1 for the average of 1 000 simulations of the across-data-type FDR, which is the average of components 2-8, excluding the global null component 1; and the misclassification rate, which is the average of all components. Our proposed methods were able to robustly control the across-data-type FDR at the prespecified *α* = 0.2 level. The separate analysis q-value failed to control the FDR. Bonferroni correction was designed to control the family-wise error rate (FWER), but we included it here to compare the misclassification rate with other methods as it is a popular error control procedure among researchers in biomedical sciences. The local FDR procedure deflated the FDR in Scenarios 3-5, which behaved similarly to the IMIX-Ind. Both methods were based on independent mixture distributions, and the reason why IMIX-Ind controlled the FDR slightly better than the local FDR procedure was that IMIX-Ind assumed a compound mixing proportion while the local FDR procedure only considered the mixing proportions for one data type at a time. For example, the mixing proportion *π*_1_ of component 1 in the IMIX-Ind is only subject to the constraint 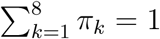, while *π*_1_ in the local FDR procedure is subject to *π*_1_ = *π*_10_*π*_20_*π*_30_, where *π*_*h*0_ is the null mixing proportion for the separate analysis of data type *h* = 1, 2, 3. This was further illustrated in Scenario 6, where the underlying correlation structures and the generating model were more complicated: the local FDR procedure failed the across-data-type FDR control while IMIX-Ind controlled it robustly. BH-FDR returned slightly inflated FDR in Scenarios 1-5, and the realized FDR increased as the correlations increased among the three data types. In Scenario 6, it failed to control the across-data-type FDR. As for the misclassification rate (Fig 1(b)), IMIX steadily achieved a lower number in all scenarios compared with commonly used methods. Our proposed methods can robustly control the across-data-type FDR and achieve a good operating characteristics under various scenarios.

**Figure 1:**
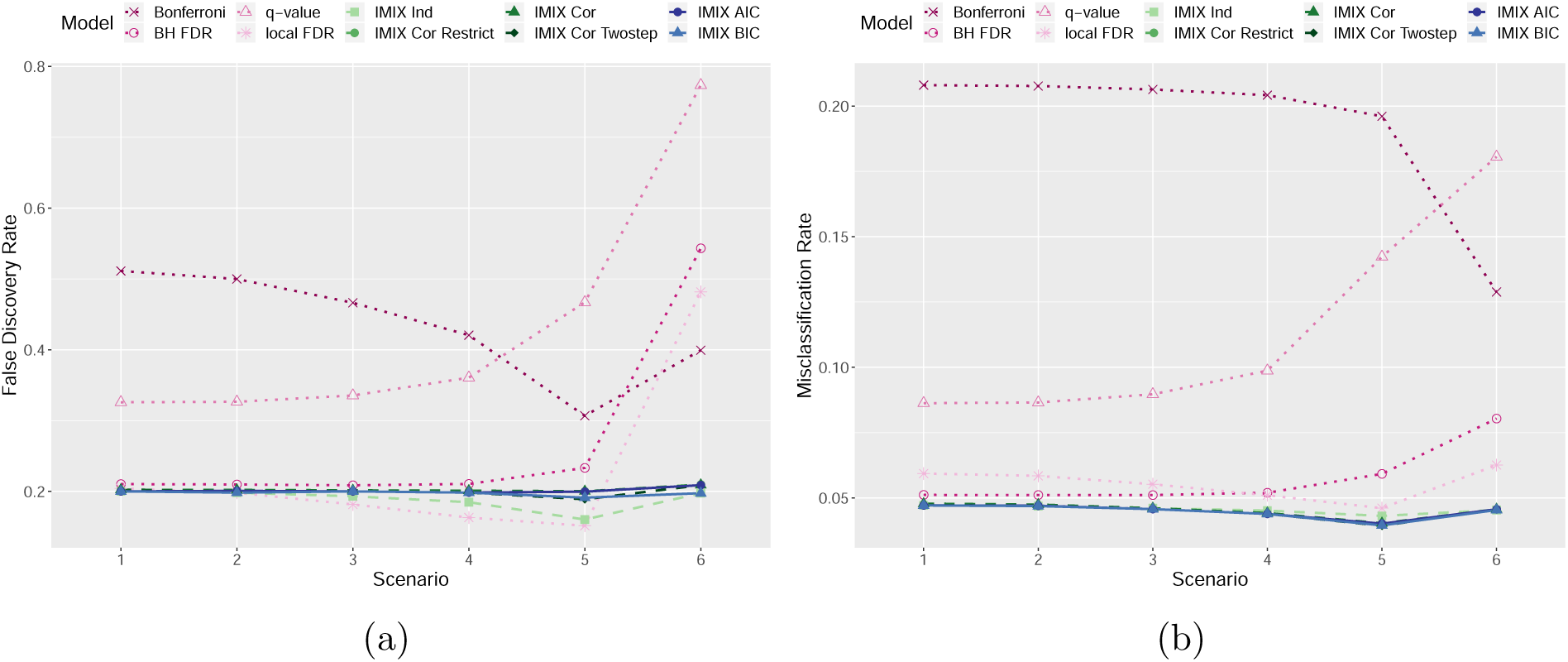
Comparison of IMIX-Ind, IMIX-Cor, IMIX-Cor-Restrict, IMIX-Cor-Twostep, IMIX-AIC, IMIX-BIC, BH-FDR, Bonferroni correction, q-value, and the local FDR procedure at *α* = 0.2: (a)across-data-type FDR control, the results are the average of 1 000 simulations of the average of components 2-8, excluding the global null component 1. (b)misclassification rate, the results are the average of 1 000 simulations of the average of all components.

In addition, we compared the computational time needed for the four IMIX models (Supplementary Material Section 2.4: table S4) using the simulation Scenario 3 assuming three data types with correlation 0.3 based on 1 000 simulations. IMIX-Ind converged the fastest within only 4.501 seconds and 67 iterations on average. Here, IMIX-Cor-Twostep achieved great computational advantages with an average of 217.379 seconds with only 42 iterations over IMIX-Cor and IMIX-Cor-Restrict, with 970.901 seconds and 417.531 seconds convergence time, 161 and 71 iterations respectively. This was processed on Intel(R) Xeon(R) CPU E5502 @ 1.87GHz with max CPU 1866 MHz and min CPU 1600 MHz.

#### 3.1.2 Model Calibration and FDR Control

Newton et al. (2004) showed that the performance of the estimated FDR based on equation

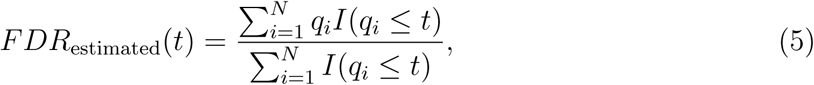

relies on how well the model fitting is. Thus, we need to assess the model calibration to ensure that IMIX framework is able to reliably control the realized FDR by the adaptive FDR procedures. We pursued this by comparing the realized and the estimated FDRs on the results fitted using IMIX from the six scenarios in simulation study 1. We compared the estimated and realized FDRs averaged across the 1 000 simulated datasets for each non-null component, and the sum up of non-nulls in comparison with the global null component 1. Fig S1-4 present the results of IMIX-Ind, IMIX-Cor-Restrict, IMIX-Cor, IMIX-Cor-Twostep in the six simulation scenarios. IMIX-Ind showed good model calibration in Scenarios 1 and 2, but as the correlation gradually increased from Scenario 3 to Scenario 5, the discrepancy between estimated FDR and realized FDR increased. In Scenario 6 where we mimicked the real data, IMIX-Ind was slightly conservative as the realized FDR was slightly smaller than the estimated FDR. IMIX-Cor and IMIX-Cor-Restrict performed similarly where the estimated and realized FDRs were coincident in scenarios 1-5. In scenario 6, component 4 and component 6 showed slightly inflated realized FDR. This was because the proportion of these two components got as small as 6.9×10^−3^ and 8.9×10^−3^. IMIX-Cor-Twostep also performed well in all scenarios except a slight shift in Scenario 5, where the correlations between data types were as high as 0.8. Since this model utilizes the estimated mean parameters from IMIX-Ind, it may have slightly affected the model calibration, but the gain in computation time was much better and can be shown in real-data based simulation Scenario 6.

In summary, the IMIX framework is rigorous and versatile to have good model calibrations under various data scenarios, which lead to a reliable and accurate FDR estimation, and thus a robust adaptive FDR control procedure.

#### 3.1.3 Model Selection

We conduct simulation study 2 to evaluate how well AIC and BIC selected the number of components in the IMIX framework. We first generated 1 000 datasets following (4) for 16 scenarios that consisted of a combination of balanced and unbalanced mixing proportions of seven and eight components (Table S1). The unbalanced mixing proportions were based on the proportions of genes in the real-data example, we used the estimated 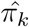 of the TCGA bladder cancer data set fitted by IMIX-Cor as the unbalanced proportions for the eight-component mixture model. The seven-component mixture model simulation simply eliminated the eighth component from the eight-component mixture model, i.e., genes that were associated with the outcome through all three data types. For each mixing proportion and number of components combination, we generated four scenarios with the mean and covariance parameters equal to the simulated parameters in simulation study 1 Scenarios 2-5. We fitted the IMIX framework without adding any constraint on the mean, assuming models of one to eight components. One reason was that it was not necessary to impose the constraints on the mean for models with fewer components; another reason was that we were only interested in the final number of components of the selected model rather than the component each gene belongs to, for the purpose of model selection.

Fig 4 shows the number of components selected by AIC/BIC after averaging 1 000 simulation study for the unbalanced seven- and eight component simulated models. AIC selected the correct number of components for the simulation study in both balanced and unbalanced settings (Fig S5(a)-(d)). BIC performed similarly (Fig 4(a)(b)(d)), but it selected a more conservative number, i.e., a more parsimonious model, under the extreme unbalanced eight-component scenario in Fig 4(c). We consider this unbalanced eight-component setting very challenging for BIC or any model selection criterion since the smallest mixing proportion was only 4‰. To further evaluate the ability of AIC and BIC to select the correct number of components when the mixing proportions were unbalanced, we conducted more simulation studies for eight-component multivariate Gaussian mixture model with varying levels of unbalanced settings as shown in Supplementary Material Section 2.3: both AIC and BIC performed well in identifying the correct number of components (Fig S6).

**Figure 2:**
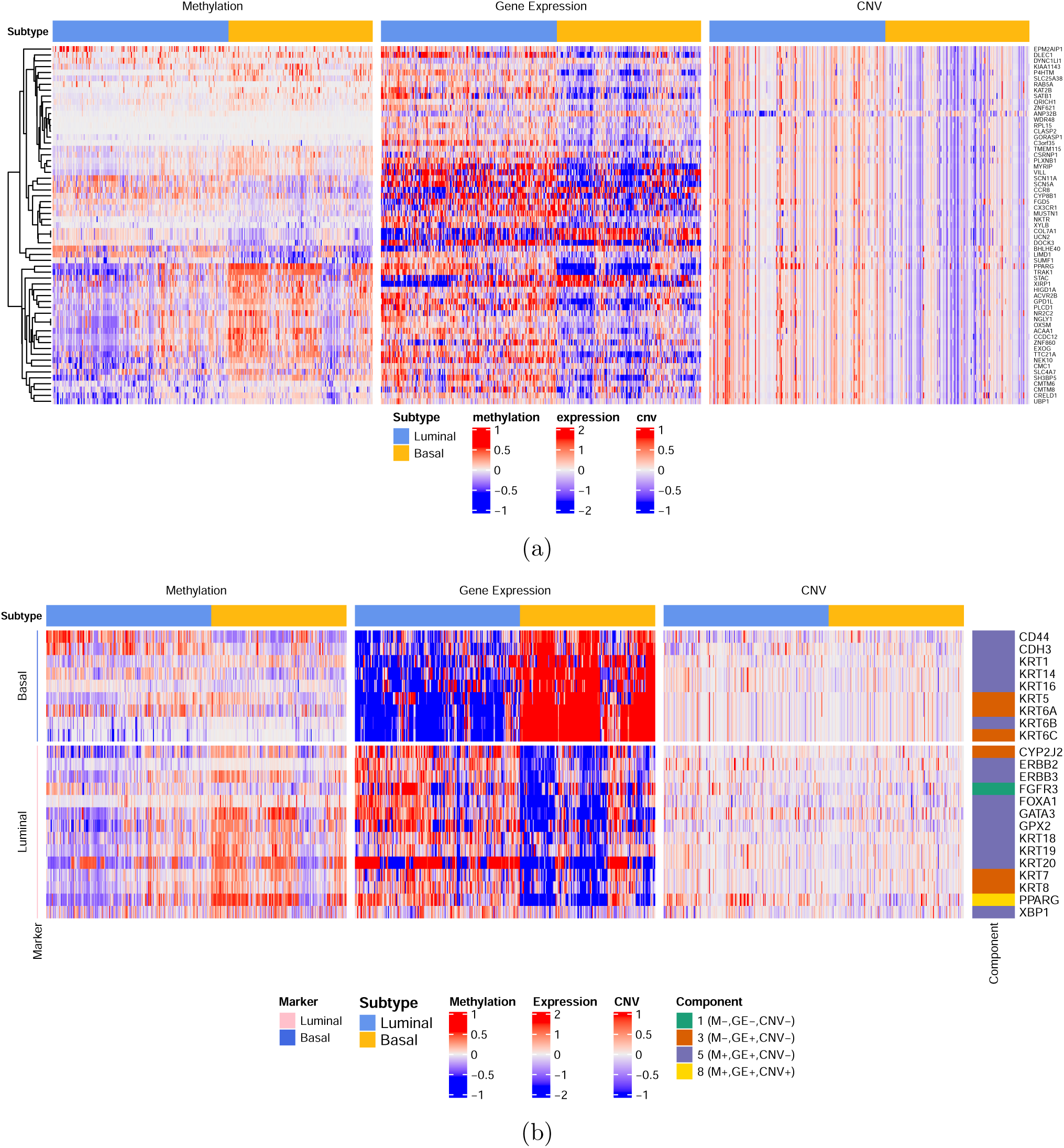
Heatmaps of genes in IMIX analysis for bladder cancer molecular subtypes in the TCGA. (a) Methylation, gene expression and copy number variation (CNV) patterns of top significant genes associated with the three data types (M+,GE+,CNV+) identified by IMIX in molecular subtypes of The Cancer Genome Atlas (TCGA) muscle-invasive bladder cancer patients, with adaptive false discovery rate (FDR) control at *α* = 0.01, estimated marginal FDR 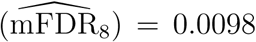. (b) Expression patterns of luminal/basal markers of TCGA bladder cancer cohort.

**Figure 3:**
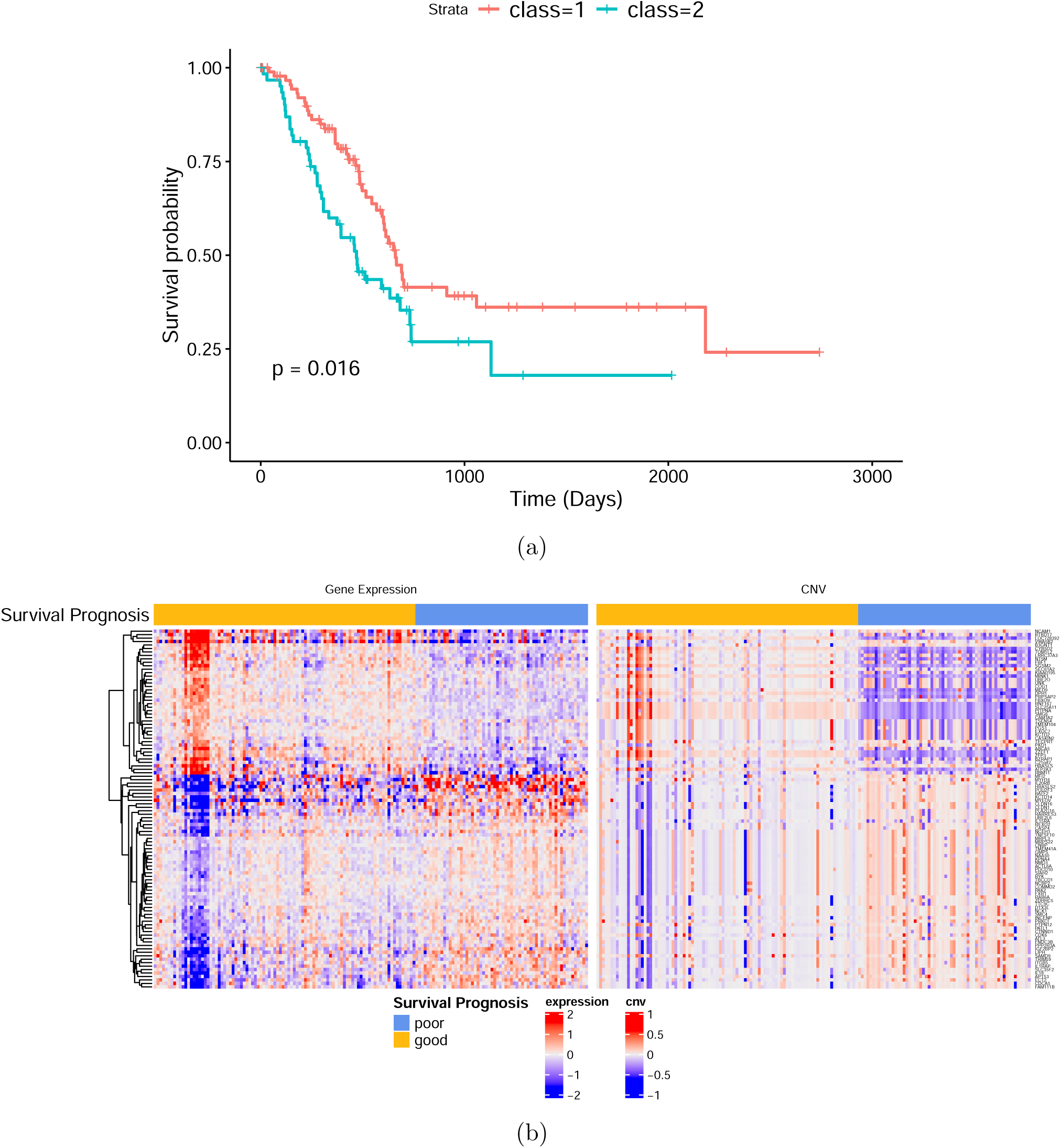
Results of IMIX analysis for pancreatic cancer prognosis in the TCGA. (a) Kaplan-Meier curves for pancreatic cancer patient survival in The Cancer Genome Atlas (TCGA). Samples were clustered based on the 104 genes identified by IMIX, with adaptive false discovery rate (FDR) control at *α* = 0.05, estimated marginal FDR 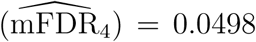. (b) Expression patterns of the 104 genes selected by IMIX that were associated with the prognosis of the TCGA pancreatic cancer patients.

**Figure 4:**
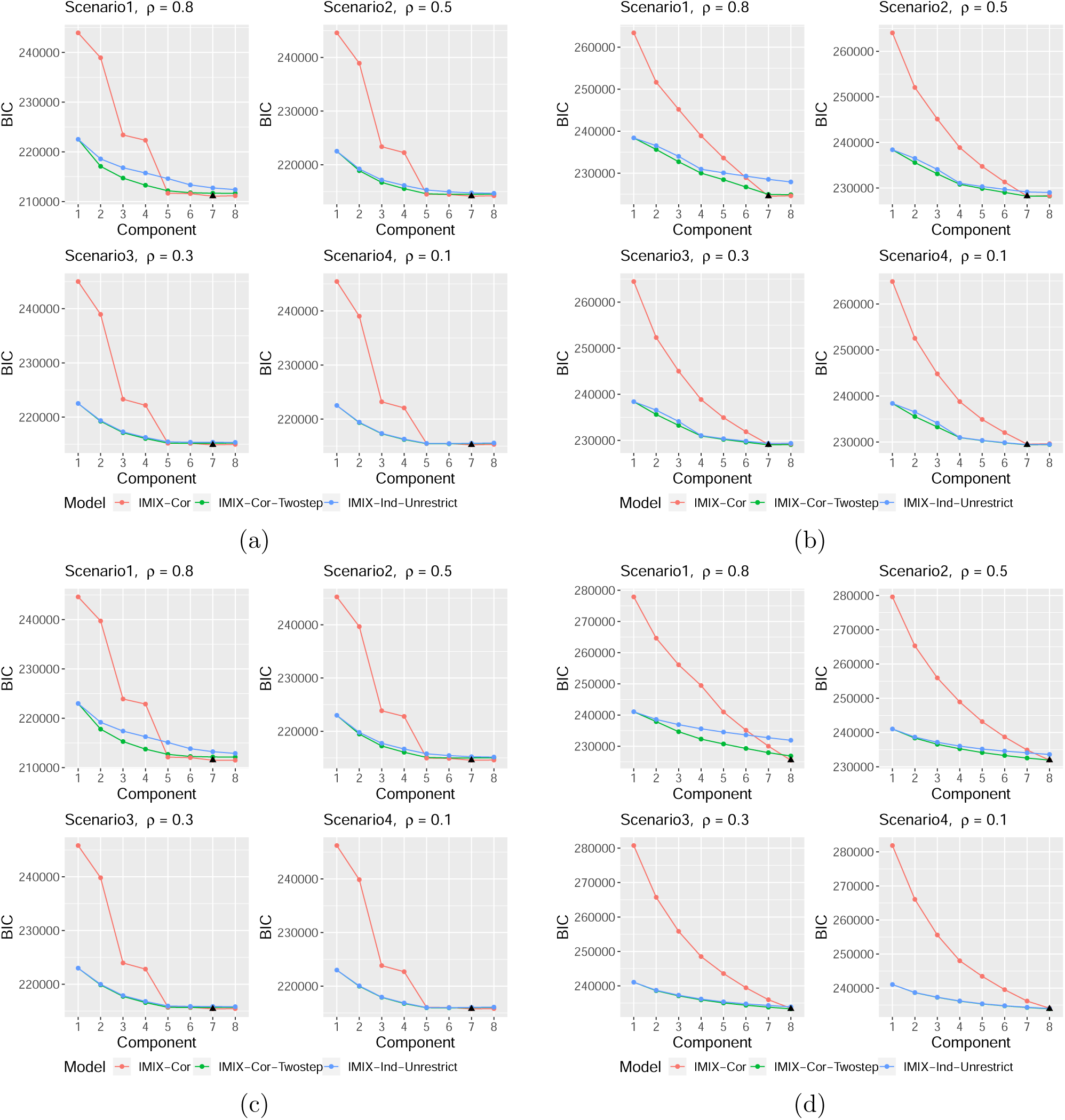
Simulation study on selecting the number of mixture components using BIC. (a)Unbalanced setting, 7-component mixture model; (b)Balanced setting, 7-component mixture model; (c)Unbalanced setting, 8-component mixture model; (d)Balanced setting, 8-component mixture model. Black triangle represents the model BIC selects. *ρ* is the correlation between data types.

Together, AIC and BIC were both reliable model selection criteria under relatively balanced mixing proportions. AIC could select up to eight components in extremely unbalanced situations; however, previous studies (McLachlan and Peel, 2004; Steele and Raftery, 2009) have shown that AIC is prone to overestimating the number of components. We consider BIC to be more stable as it takes into account the number of genes in the penalty term, which can be as large as thousands under whole genome setting.

### 3.2 Real Data Applications

To demonstrate the proposed IMIX framework’s versatility and efficiency in different disease outcomes, we applied our method to a binary outcome, the luminal and basal molecular subtypes of muscle-invasive bladder cancer, as well as a survival outcome for the prognosis of pancreatic cancer in the TCGA dataset.

#### 3.2.1 Molecular Subtypes of Bladder Cancer in the TCGA

Previous studies in bladder cancer identified molecular signatures associated with the pathological and clinical outcomes (Choi et al., 2014; Guo et al., 2019); in particular, those molecular subtypes have important implications for prognostication and treatment. Twenty-three gene expression markers have been reported to play a major role in these molecular subtypes. Here we applied IMIX to jointly analyze DNA methylation, gene expression, and CNV to investigate: (1) whether those gene expression markers also demonstrated difference at the DNA methylation and CNV levels, and (2) whether there were other genes associated with the molecular subtyping through any of the three data types. We analyzed the TCGA bladder cancer patient cohort that was profiled by three genomic platforms, DNA methylation, mRNA gene expression, and CNV. After quality control (Supplementary Materials, Section 3.1), we separately analyzed 373 DNA methylation samples, 391 RNA-Seq samples, and 387 CNV samples with *N* = 15 672 genes with respect to the molecular subtypes adjusting for the clinical covariates, including age, sex, race, smoking status and pathologic stage. We applied IMIX, Bonferroni correction, and BH-FDR to the final summary statistics/Z-scores obtained from the association tests of individual-level data. The nominal error control level of Bonferroni correction and BH-FDR for separate analysis was set at *α* = 0.05 and that of IMIX for integrative analysis was at *α* = 0.2. We used IMIX-BIC to perform model selection, with the optimal model selected as IMIX-Cor-Twostep and the best number of components as eight based on BIC values. Table 1 shows the point estimates and 95% bootstrap-based confidence intervals (*B* = 1 000) (McLachlan and Peel, 2004) for the parameters in the correlation matrices between DNA methylation, gene expression, and CNV. DNA methylation and gene expression were correlated for all components that involved non-null genes through at least one of these two data types: they were component 2 (M+,E−,CNV−), component 3 (M−,E+,CNV−), component 5 (M+,E+,CNV−), and component 8 (M+,E+,CNV+). In particular, component 5 and component 8 showed moderate correlations between DNA methylation and gene expression where genes showed significant associations through both data types (M+ and E+). Another interesting finding was that the two data types DNA methylation and gene expression were correlated when genes were associated with the outcome through only one datatype as reflected in component 2 (M+ and E−) and component 3 (M− and E+). This also held for the correlations between gene expression and CNV in component 1 (E−,CNV−) and component 3 (E+,CNV−). These results further supported the IMIX model assumptions that the data types were correlated, both under the alternative and the null hypothesis, thus reinforcing that the IMIX method was effective by assuming multivariate distributions in all components instead of the commonly adopted conditional independence under the null hypothesis.

**Table 1:** Estimated correlations between the transformed z scores (from p value) of DNA methylation (M) and gene expression (E), DNA methylation and CNV, gene expression and CNV with 95% bootstrap-based confidence intervals (B = 1 000) for TCGA bladder cancer data integration analysis by IMIX-BIC.

| Component | M & E | M & CNV | E & CNV |
| --- | --- | --- | --- |
| 1 (M-,E-,CNV-) | 0.015 (-0.039,0.072) | **0.091 (0.037,0.14)** | -0.016 (-0.070,0.043) |
| 2 (M+,E-,CNV-) | **0.070 (0.014,0.12)** | -0.012 (-0.060,0.031) | 0.0049 (-0.064,0.075) |
| 3 (M-,E+,CNV-) | **0.14 (0.051,0.23)** | **0.090 (0.016,0.17)** | **-0.10 (-0.19,-0.022)** |
| 4 (M-,E-,CNV+) | -0.099 (-0.59,0.40) | 0.35 (-0.085,0.73) | -0.059 (-0.53,0.36) |
| 5 (M+,E+,CNV-) | **0.25 (0.20,0.31)** | -0.037 (-0.082,0.013) | 0.038 (-0.023,0.097) |
| 6 (M+,E-,CNV+) | 0.038 (-0.22,0.25) | -0.037 (-0.27,0.22) | -0.088 (-0.40,0.26) |
| 7 (M-,E+,CNV+) | 0.21 (-0.16,0.56) | 0.13 (-0.27,0.49) | 0.12 (-0.29,0.50) |
| 8 (M+,E+,CNV+) | **0.19 (0.021,0.38)** | 0.10 (-0.049,0.26) | 0.080 (-0.13,0.28) |

We compared the number of genes discovered in component 8 using BH-FDR, Bonferroni correction, and our method (Fig S7(a)). The genes that were detected by Bonferroni correction were identified by both our method and BH-FDR. The genes detected by IMIX had an overlap of 146 genes with the BH-FDR and included 116 new genes not discovered by either BH-FDR or Bonferroni correction. The estimated 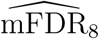 of IMIX was 0.1995, close to the prespecified across-data-type FDR control level alpha=0.2. Through simulation studies in Section 3.1.1, our method was more effective in controlling the across-data-type FDR compared with other methods. We also showed the levels of DNA methylation, gene expression, and CNV for the significant genes in component 8, i.e., genes that were associated with all the three data types in Fig 2(a); for the purpose of illustration, we only included the 61 significant genes after adaptive FDR control at *α* = 0.01. We conducted Ingenuity Pathway Analysis (IPA, Ingenuity Systems (www.ingenuity.com)) on the 61 significant genes in component 8 at *α* = 0.01. The results showed strong peroxisome proliferator activator receptor (PPAR) pathway activation (Fig S9) in luminal samples. This pathway was previously reported by Choi et al. (2014), who first proposed the molecular subtypes of muscle-invarsive bladder cancer, that PPAR*α* and PPAR*γ* activation played essential roles in regulating gene expression signature for the luminal subtype. Specifically, they exposed the PPAR*γ*-selective agonist rosiglitazone in two bladder cancer cell lines and further confirmed that rosiglitazone activated PPAR pathway and enriched gene signatures in primary luminal samples. Furthermore, we estimated the causal relationships between DNA methylation, gene expression, and CNV of the 61 genes in component 8 by applying Bayesian networks (Scutari, 2017) with the target nominal type I error rate at 0.01. Among which, 51 genes showed significant dependent structures between the three data types. The directed acyclic graphs (DAGs) based on conditional independence tests with a restriction of causal direction from CNV to E showed six different patterns of causal structures (Supplementary Materials, Section 3.2).

We present the levels of the luminal/basal markers for DNA methylation, gene expression, and CNV in Fig 2(b). Among the 23 markers, we found that six, fifteen, and one gene belonged to component 3 (M-,E+,CNV-), component 5 (M+,E+,CNV-), and component 8 (M+,E+,CNV+), respectively. In particular, PPARG belonged to component 8, i.e, associated with the subtypes via all three molecular mechanisms. Bayesian networks further confirmed that PPARG had a full model with dependence structures of CNV*→*E, E−M, M−CNV (Fig S8(1)). This gene was reported to be one of the driver genes for the basal/luminal differentiation. As expected, PPARG showed higher gene expression level in the luminal samples than in the basal samples; furthermore, we discovered a concordant significant differential pattern in the methylation and CNV levels that have not been previously reported.

In summary, our analysis revealed that the luminal/basal markers demonstrated substantial differences in at least two data types (Fig 2(b)). By applying the IMIX framework, we successfully discovered novel genes that were associated with the molecular subtypes through all three data types (Fig 2(a)) and confirmed the PPAR/RXR activation canonical pathway that was previously reported to play a central role in luminal/basal differentiation (Choi et al., 2014).

#### 3.2.2 Prognosis of Pancreatic Cancer in the TCGA

We further applied IMIX to a survival outcome to investigate the relationships between the prognosis of pancreatic cancer patients and two genomic datasets, gene expression and CNV in the TCGA. After quality control (Supplementary Materials, Section 3.1), we first applied the Cox proportional hazards model to each of the 15 472 genes respectively on 157 RNA-Seq samples and 161 CNV samples adjusting for age, gender, and smoking status. Next, we fitted IMIX, BH-FDR, and Bonferroni corrections on the summary statistics. After model selection based on BIC, IMIX-Cor-Twostep fitted the best. Table 2 shows the point estimates and 95% bootstrap-based confidence intervals (*B* = 1 000) of the parameters in the correlation matrices between gene expression and CNV. In component 4 (E+,CNV+), where the detected genes were significantly associated with survival outcomes through both gene expression and CNV, the correlation between gene expression and CNV was 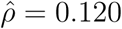 with 95% confidence interval (0.071, 0.18). To assess the effect of the detected 104 genes in component 4 at *α* = 0.05, we used iCluster (Shen et al., 2009) to group the patients based on gene expression and CNV data into two classes, here we only applied the 104 genes detected by IMIX with no feature selection in the clustering process. Fig 3(a) shows the Kaplan-Meier(KM) curve of the overall survival of the pancreatic cancer patients. The log-rank test resulted in *p* = 0.016, and the Cox model adjusting for patient pathologic stages resulted in *p* = 0.04. Furthermore, we also clustered the patients using the 991 genes discovered at adaptive FDR *α* = 0.2, all patients but one were grouped into the same clusters as using the 104 genes at *α* = 0.05; the KM curve returned the same results. This indicates that IMIX was able to capture the most important features and a controlled number of false discovered genes at *α* = 0.05. Fig 3(b) shows the gene expression and CNV levels of the identified 104 genes associated with the pancreatic cancer prognosis. The gene expression and CNV were positively correlated as shown in the heatmap. We compared the results of IMIX, Bonferonni correction, and BH-FDR for component 4, i.e., genes that were associated with the survival outcomes through both gene expression and CNV. Bonferroni correction was not able to discover any significant genes at the nominal level *α* = 0.05. IMIX detected 104 genes at *α* = 0.05 with an estimated 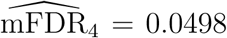. BH-FDR detected 271 genes at *α* = 0.05. IMIX identified fewer genes but it captured the important features as evidenced by the KM analysis/log-rank test with a controlled across-data-type FDR compared with BH-FDR. We showed in Section 3.1.1 that BH-FDR failed to control for FDR under the data integration settings. In addition, IMIX detected 991 genes at *α* = 0.2 with an estimated 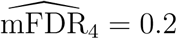; and the 271 genes detected by BH-FDR (*α* = 0.05) were all included in the genes discovered by our method as shown in the Venn diagram (Fig S7(b)).

**Table 2:** Estimated correlations between the transformed z scores (from p value) of gene expression (E) and CNV with 95% bootstrap-based confidence intervals (B = 1 000) for TCGA pancreatic cancer data integration analysis by IMIX-BIC.

| Component | E & CNV |
| --- | --- |
| 1 (E-,CNV-) | 0.040 (-0.047,0.13) |
| 2 (E+,CNV-) | 0.11 (-0.012,0.21) |
| 3 (E-,CNV+) | 0.016 (-0.019,0.049) |
| 4 (E+,CNV+) | **0.12 (0.071,0.18)** |

## 4 Discussion

We have proposed IMIX, a multivariate mixture model framework based on summary statistics for integrative genomic association analysis. Our model incorporates the correlation structures between different genomic datasets by assuming multivariate Gaussian mixture distribution of the Z scores (transformed from p-values) from regression analysis of individual-level data. The IMIX framework includes four models: IMIX-Cor, IMIX-Ind, IMIX-Cor-Restrict, and IMIX-Cor-Twostep, each of which best captures a specific type of data correlation structure arising from various data analysis problems. IMIX selects the optimal model based on AIC/BIC values among the four models. In addition, it features simultaneous model selection for the number of underlying latent states/components of the optimal mixture model with a specific correlation structure. We utilize the EM algorithm in parameter estimation, and the mixture model naturally produces the local FDR for each gene, which is easily derived from the posterior probability. Our model features an adaptive procedure to control the across-data-type FDR, where we take into account both the multiple testing of the gene, and the multiple data types under an integrative analysis setting. To our knowledge, we are the first to demonstrate such error-control property of integrative genomic models.

Our applications to the two TCGA data sets demonstrate that different genomic data types, such as DNA methylation, mRNA gene expression, and CNVs, can be correlated in both null and non-null genes, as shown in the bootstrap-based confidence intervals (Table 1 and 2). Therefore, it is necessary to consider the inter-source correlations of multiple datasets in integrative analysis. Based on simulation studies under various settings of correlation structures, including the one based on TCGA bladder cancer dataset, IMIX controlled the FDR precisely and yielded better statistical power compared with the independent separate analysis models, including BH-FDR, Bonferroni correction, q-value, and the local FDR procedure.

An advantage of our proposed method using summary statistics is that our method does not require the use of a common set of samples, which relaxes the conditions of previously published methods for the integration of multiple omics data (Richardson et al., 2016). In addition, IMIX is able to model summary statistics from independent or partially overlapped cohorts, as illustrated in the TCGA data examples in Section 3.2. The implementation of IMIX employs the EM algorithm, which in general converges fast, leading to great computational efficiency. One unique feature of IMIX is its constraint on the mixture component means, which is not only more biologically plausible than unconstrainted means, but has also ensured model identifiability, a common challenge in mixture models. Another unique feature of IMIX is model selection: we let the data decide the number of mixture components and the correlation structure based on AIC or BIC.

We compare our work with a recently proposed method called Primo for quantitative trait loci (QTLs) mapping based on genome-wide association study (GWAS) summary statistics (Gleason et al., 2019). The two methods share a similar concept in utilizing the mixture model in data integration; however, there are several key differences. The main difference is how we approach the parameter estimations. Primo estimates the mixture model parameters by first assuming conditional independence between the data types and estimating the marginal null and alternative distributions for each data type with a fixed proportion of non-null tests, and then approximating the correlation matrices under certain assumptions. In contrast, IMIX directly estimates the multivariate mixture model parameters, including means, covariance matrices and mixing proportions, using the EM algorithm, which allows examining and relaxing the conditional independence assumption. Furthermore, IMIX accommodates simultaneous model selection for both number of mixture components and the correlation structure, which is absent in Primo.

IMIX is a useful and versatile tool that can study various types of outcomes, including continuous, binary, and time-to-event outcomes in integrative genomic analysis. We have applied IMIX to two types of problems, the survival prognosis of pancreatic cancer and the luminal/basal molecular subtypes of bladder cancer, both providing novel biological insights. IMIX framework is not only applicable to cancer genomics, but also to other complex diseases and traits as afforded by ongoing large-scale multiple-omics projects, such as the NIH Trans-Omics for Precision Medicine (TOPMed) project (Brody et al., 2017) and the Cohorts for Heart and Aging Research in Genomic Epidemiology (CHARGE) Consortium (Psaty et al., 2009), consisting of over 100,000 deeply phenotyped and sequenced individuals with multiple types of omics data, such as transcriptomic, epigenomic, metabolic, proteomic, and whole-genome sequencing data. This work, therefore, has a wide range of application potential to provide novel biological insights into disease mechanisms.We have implemented the integration model for two and three genomics data types in the simulation studies and data applications, which could be further generalized to four and more data types in the multivariate mixture model framework.We leave the details of this potential extension for future research. While we have relaxed the conditional independence assumptions for the data types in IMIX, we could further extend our method by assessing the correlations between genes within each data type, which is another important direction for future work.

We have implemented the proposed method in an R package “IMIX”, which is available at https://github.com/ziqiaow/IMIX and will be posted to R/CRAN.

## Acknowledgements

This research was supported by the National Institutes of Health (NIH) grants R01HL116720 and R01CA169122; P.W. was supported by NIH grant P50CA091846. The results shown here are in whole or part based upon data generated by the TCGA Research Network: https://www.cancer.gov/tcga. We thank Ms. Sarah Bronson in Scientific Publications, Research Medical Library, The University of Texas MD Anderson Cancer Center, for editorial assistance. The authors declare no conflict of interest.

