## Supplementary Materials for "IMIX: A multivariate mixture model approach to integrative analysis of multiple types of omics data"

Ziqiao Wang[1,2] and Peng Wei[1,*]

[1]Department of Biostatistics, The University of Texas MD Anderson Cancer Center, Houston, Texas, USA
[2]The University of Texas MD Anderson Cancer Center UTHealth Graduate School of Biomedical Sciences, Houston, Texas, USA

#### 1 Details on the EM algorithm

##### 1.1 IMIX-Ind

To infer whether gene i is associated with data type h, we use the posterior probability

$$\begin{aligned} Pr(z_{ik} = 1|X_{i1}, X_{i2}, X_{i3}, \boldsymbol{\theta}) =& \frac{\pi_k f_k(X_{i1}, X_{i2}, X_{i3})}{\sum_{j=1}^K \pi_j f_j(X_{i1}, X_{i2}, X_{i3})} \\ =& \frac{\pi_k f_{k1}(X_{i1}; \theta_{k1}) f_{k2}(X_{i2}; \theta_{k2}) f_{k3}(X_{i3}; \theta_{k3})}{f(X_{i1}, X_{i2}, X_{i3})}. \end{aligned}$$

Notice that here we assume $K = 8$. For each scenario, it corresponds as:

K=1: (0,0,0); K=2: (1,0,0); K=3: (0,1,0); K=4: (0,0,1); K=5: (1,1,0); K=6: (1,0,1); K=7: (0,1,1); K=8: (1,1,1).

*Correspondence should be addressed to: Peng Wei, PhD, 1400 Pressler St, Unit 1411, Department of Biostatistics, The University of Texas MD Anderson Cancer Center, Houston, TX 77030, USA;

We assume $f_{kh} = \phi(.; \mu_{kh}, \sigma_{kh})$, a normal probability density function with mean $\mu_{kh}$ and variance $\sigma_{kh}^2$, $k = 1, \cdots, 8; h = 1, 2, 3$. Thus we have

$$\begin{aligned}
f(X_{i1}, X_{i2}, X_{i3}) =& \sum_{k=1}^{K} \pi_k f_k(x_1, x_2, x_3) \\
=& \sum_{k=1}^{K} \pi_k f_{k1}(x_1; \theta_{k1}) f_{k2}(x_2; \theta_{k2}) f_{k3}(x_3; \theta_{k3}) \\
=& \pi_1 f(x_1; \mu_{10}, \sigma_{10}) f(x_2; \mu_{20}, \sigma_{20}) f(x_3; \mu_{30}, \sigma_{30}) \\
&+ \pi_2 f(x_1; \mu_{11}, \sigma_{11}) f(x_2; \mu_{20}, \sigma_{20}) f(x_3; \mu_{30}, \sigma_{30}) \\
&+ \pi_3 f(x_1; \mu_{10}, \sigma_{10}) f(x_2; \mu_{21}, \sigma_{21}) f(x_3; \mu_{30}, \sigma_{30}) \\
&+ \pi_4 f(x_1; \mu_{10}, \sigma_{10}) f(x_2; \mu_{20}, \sigma_{20}) f(x_3; \mu_{31}, \sigma_{31}) \\
&+ \pi_5 f(x_1; \mu_{11}, \sigma_{11}) f(x_2; \mu_{21}, \sigma_{21}) f(x_3; \mu_{30}, \sigma_{30}) \\
&+ \pi_6 f(x_1; \mu_{11}, \sigma_{11}) f(x_2; \mu_{20}, \sigma_{20}) f(x_3; \mu_{31}, \sigma_{31}) \\
&+ \pi_7 f(x_1; \mu_{10}, \sigma_{10}) f(x_2; \mu_{21}, \sigma_{21}) f(x_3; \mu_{31}, \sigma_{31}) \\
&+ \pi_8 f(x_1; \mu_{11}, \sigma_{11}) f(x_2; \mu_{21}, \sigma_{21}) f(x_3; \mu_{31}, \sigma_{31}).
\end{aligned}$$

$$\begin{aligned}
&\mu_{11} = \mu_{31} = \mu_{41} = \mu_{71} \overset{\triangle}{=} \mu_{10}, \sigma_{11} = \sigma_{31} = \sigma_{41} = \sigma_{71} \overset{\triangle}{=} \sigma_{10} \\
&\mu_{12} = \mu_{22} = \mu_{42} = \mu_{62} \overset{\triangle}{=} \mu_{20}, \sigma_{12} = \sigma_{22} = \sigma_{42} = \sigma_{62} \overset{\triangle}{=} \sigma_{20} \\
&\mu_{13} = \mu_{23} = \mu_{33} = \mu_{53} \overset{\triangle}{=} \mu_{30}, \sigma_{13} = \sigma_{23} = \sigma_{33} = \sigma_{43} \overset{\triangle}{=} \sigma_{30} \\
&\mu_{21} = \mu_{51} = \mu_{61} = \mu_{81} \overset{\triangle}{=} \mu_{11}, \sigma_{21} = \sigma_{51} = \sigma_{61} = \sigma_{81} \overset{\triangle}{=} \sigma_{11} \\
&\mu_{32} = \mu_{52} = \mu_{72} = \mu_{82} \overset{\triangle}{=} \mu_{21}, \sigma_{32} = \sigma_{52} = \sigma_{72} = \sigma_{82} \overset{\triangle}{=} \sigma_{21} \\
&\mu_{43} = \mu_{63} = \mu_{73} = \mu_{83} \overset{\triangle}{=} \mu_{31}, \sigma_{43} = \sigma_{63} = \sigma_{73} = \sigma_{83} \overset{\triangle}{=} \sigma_{31}.
\end{aligned}$$

For the EM algorithm, the complete log-likelihood is

$$\log L_c = \sum_{i=1}^{N}\sum_{k=1}^{K} z_{ik} \log \pi_k f_k(X_{i1}, X_{i2}, X_{i3}).$$

The E-step is to calculate the conditional expectation

$$Q = E(\log L_c | data) = \sum_{i=1}^{N}\sum_{k=1}^{K} \tau(z_{ik}) \log \pi_k f_k(X_{i1}, X_{i2}, X_{i3}),$$

where $\tau(z_{ik}) = Pr(z_{ik} = 1 | X_{i1}, X_{i2}, X_{i3})$. The M-step maximized the above Q with respect to the unknown parameters:

**E step:** Compute $\tau(z_{ik})$ with current parameter $\boldsymbol{\theta}^{(m)} = \{\pi_k^{(m)}, \mu_{kh}^{(m)}, {\sigma_{kh}^2}^{(m)}\}$ at each iteration m:

$$\tau(z_{ik}) = Pr(z_{ik} = 1 | X_{i1}, X_{i2}, X_{i3}) = \frac{\pi_k^{(m)} f_k^{(m)}(X_{i1}, X_{i2}, X_{i3})}{\sum_{j=1}^{K} \pi_j^{(m)} f_j^{(m)}(X_{i1}, X_{i2}, X_{i3})},$$

where $f_k^{(m)}(X_{i1}, X_{i2}, X_{i3}) = \prod_{h=1}^{3} \phi(X_{ih}; \mu_{kh}^{(m)}, \sigma_{kh}^{(m)})$.

**M step:** Update $\boldsymbol{\theta}^{(m)}$ and replace by $\boldsymbol{\theta}^{(m+1)}$:

$$
\begin{aligned}
\pi_k^{(m+1)} &= \frac{\sum_{i=1}^{N} \tau(z_{ik})}{N}, \\
\mu_{10}^{(m+1)} &= \frac{\sum_{i=1}^{N} X_{i1}(\tau(z_{i1}) + \tau(z_{i3}) + \tau(z_{i4}) + \tau(z_{i7}))}{\sum_{i=1}^{N}(\tau(z_{i1}) + \tau(z_{i3}) + \tau(z_{i4}) + \tau(z_{i7}))}, \\
\mu_{20}^{(m+1)} &= \frac{\sum_{i=1}^{N} X_{i2}(\tau(z_{i1}) + \tau(z_{i2}) + \tau(z_{i4}) + \tau(z_{i6}))}{\sum_{i=1}^{N}(\tau(z_{i1}) + \tau(z_{i2}) + \tau(z_{i4}) + \tau(z_{i6}))}, \\
\mu_{30}^{(m+1)} &= \frac{\sum_{i=1}^{N} X_{i3}(\tau(z_{i1}) + \tau(z_{i2}) + \tau(z_{i3}) + \tau(z_{i5}))}{\sum_{i=1}^{N}(\tau(z_{i1}) + \tau(z_{i2}) + \tau(z_{i3}) + \tau(z_{i5}))}, \\
\mu_{11}^{(m+1)} &= \frac{\sum_{i=1}^{N} X_{i1}(\tau(z_{i2}) + \tau(z_{i5}) + \tau(z_{i6}) + \tau(z_{i8}))}{\sum_{i=1}^{N}(\tau(z_{i2}) + \tau(z_{i5}) + \tau(z_{i6}) + \tau(z_{i8}))}, \\
\mu_{21}^{(m+1)} &= \frac{\sum_{i=1}^{N} X_{i2}(\tau(z_{i3}) + \tau(z_{i5}) + \tau(z_{i7}) + \tau(z_{i8}))}{\sum_{i=1}^{N}(\tau(z_{i3}) + \tau(z_{i5}) + \tau(z_{i7}) + \tau(z_{i8}))}, \\
\mu_{31}^{(m+1)} &= \frac{\sum_{i=1}^{N} X_{i3}(\tau(z_{i4}) + \tau(z_{i6}) + \tau(z_{i7}) + \tau(z_{i8}))}{\sum_{i=1}^{N}(\tau(z_{i4}) + \tau(z_{i6}) + \tau(z_{i7}) + \tau(z_{i8}))}, \\
\sigma_{10}^{2\ (m+1)} &= \frac{\sum_{i=1}^{N}(X_{i1} - \mu_{10}^{(m+1)})^2(\tau(z_{i1}) + \tau(z_{i3}) + \tau(z_{i4}) + \tau(z_{i7}))}{\sum_{i=1}^{N}(\tau(z_{i1}) + \tau(z_{i3}) + \tau(z_{i4}) + \tau(z_{i7}))}, \\
\sigma_{20}^{2\ (m+1)} &= \frac{\sum_{i=1}^{N}(X_{i2} - \mu_{20}^{(m+1)})^2(\tau(z_{i1}) + \tau(z_{i2}) + \tau(z_{i4}) + \tau(z_{i6}))}{\sum_{i=1}^{N}(\tau(z_{i1}) + \tau(z_{i2}) + \tau(z_{i4}) + \tau(z_{i6}))}, \\
\sigma_{30}^{2\ (m+1)} &= \frac{\sum_{i=1}^{N}(X_{i3} - \mu_{30}^{(m+1)})^2(\tau(z_{i1}) + \tau(z_{i2}) + \tau(z_{i3}) + \tau(z_{i5}))}{\sum_{i=1}^{N}(\tau(z_{i1}) + \tau(z_{i2}) + \tau(z_{i3}) + \tau(z_{i5}))}, \\
\sigma_{11}^{2\ (m+1)} &= \frac{\sum_{i=1}^{N}(X_{i1} - \mu_{11}^{(m+1)})^2(\tau(z_{i2}) + \tau(z_{i5}) + \tau(z_{i6}) + \tau(z_{i8}))}{\sum_{i=1}^{N}(\tau(z_{i2}) + \tau(z_{i5}) + \tau(z_{i6}) + \tau(z_{i8}))}, \\
\sigma_{21}^{2\ (m+1)} &= \frac{\sum_{i=1}^{N}(X_{i2} - \mu_{21}^{(m+1)})^2(\tau(z_{i3}) + \tau(z_{i5}) + \tau(z_{i7}) + \tau(z_{i8}))}{\sum_{i=1}^{N}(\tau(z_{i3}) + \tau(z_{i5}) + \tau(z_{i7}) + \tau(z_{i8}))}, \\
\sigma_{31}^{2\ (m+1)} &= \frac{\sum_{i=1}^{N}(X_{i3} - \mu_{31}^{(m+1)})^2(\tau(z_{i4}) + \tau(z_{i6}) + \tau(z_{i7}) + \tau(z_{i8}))}{\sum_{i=1}^{N}(\tau(z_{i4}) + \tau(z_{i6}) + \tau(z_{i7}) + \tau(z_{i8}))}.
\end{aligned}
$$

Repeat the above iterations until convergence.

#### 1.2 IMIX-Cor

We assume that the three data sources can be summarized as $(X_{i1}, X_{i2}, X_{i3})$ for each gene $i, i = 1, \cdots, N$, data type $h = 1, 2, 3$: $X_{ih} = \Phi^{-1}(p_{ih})$, $p_{ih}$ is the p-values. We assume that $(X_{i1}, X_{i2}, X_{i3})$ comes from a mixture distribution with K mixture components:

$$f(x_1, x_2, x_3) = \sum_{k=1}^{K} \pi_k f_k(x_1, x_2, x_3).$$

To infer whether gene i is associated with data type h, we use the posterior probability

$$Pr(z_{ik} = 1 | X_{i1}, X_{i2}, X_{i3}, \boldsymbol{\theta}) = \frac{\pi_k f_k(X_{i1}, X_{i2}, X_{i3})}{\sum_{j=1}^{K} \pi_j f_j(X_{i1}, X_{i2}, X_{i3})}.$$

Notice that here we assume $K = 8$. For each scenario, it corresponds as:
K=1: (0,0,0); K=2: (1,0,0); K=3: (0,1,0); K=4: (0,0,1); K=5: (1,1,0); K=6: (1,0,1); K=7: (0,1,1); K=8: (1,1,1).

We assume $f_k = N(\boldsymbol{\mu_k, \Sigma_k})$ with mixing proportion $\pi_k, k = 1, \cdots, 8$. Here we add a constrain on the $\boldsymbol{\mu}_k$:

$$\boldsymbol{\mu_1} = (\mu_{10}, \mu_{20}, \mu_{30}); \boldsymbol{\mu_2} = (\mu_{11}, \mu_{20}, \mu_{30});$$
$$\boldsymbol{\mu_3} = (\mu_{10}, \mu_{21}, \mu_{30}); \boldsymbol{\mu_4} = (\mu_{10}, \mu_{20}, \mu_{31});$$
$$\boldsymbol{\mu_5} = (\mu_{11}, \mu_{21}, \mu_{30}); \boldsymbol{\mu_6} = (\mu_{11}, \mu_{20}, \mu_{31});$$
$$\boldsymbol{\mu_7} = (\mu_{10}, \mu_{21}, \mu_{31}); \boldsymbol{\mu_8} = (\mu_{11}, \mu_{21}, \mu_{31}).$$

The complete log-likelihood is

$$\log L_c = \sum_{i=1}^{N} \sum_{k=1}^{K} z_{ik} \log \pi_k f_k(X_{i1}, X_{i2}, X_{i3}).$$

The E-step is to calculate the conditional expectation

$$Q = E(\log L_c | data) = \sum_{i=1}^{N} \sum_{k=1}^{K} \tau(z_{ik}) \log \pi_k f_k(X_{i1}, X_{i2}, X_{i3}).$$

where $\tau(z_{ik}) = Pr(z_{ik} = 1 | X_{i1}, X_{i2}, X_{i3})$. The M-step maximized the above Q with respect to the unknown parameters, for the unconstrained model:

**E step:** Compute $\tau(z_{ik})$ with current parameter $\boldsymbol{\theta}^{(m)} = \{\pi_k^{(m)}, \boldsymbol{\mu_k}^{(m)}, \boldsymbol{\Sigma_k}^{(m)}\}$ at each iteration m:

$$\tau(z_{ik}) = Pr(z_{ik} = 1 | X_{i1}, X_{i2}, X_{i3}) = \frac{\pi_k^{(m)} f_k^{(m)}(X_{i1}, X_{i2}, X_{i3})}{\sum_{j=1}^{K} \pi_j^{(m)} f_j^{(m)}(X_{i1}, X_{i2}, X_{i3})},$$

where $f_k^{(m)}(X_{i1}, X_{i2}, X_{i3}) = \mathcal{N}(\boldsymbol{X_i} | \boldsymbol{\mu_k}^{(m)}, \boldsymbol{\Sigma_k}^{(m)})$.

**M step:** Update $\boldsymbol{\theta}^{(m)}$ and replace by $\boldsymbol{\theta}^{(m+1)}$:

$$\begin{aligned}
\pi_k^{(m+1)} &= \frac{\sum_{i=1}^{N} \tau(z_{ik})}{N}, \\
\boldsymbol{\mu}_k^{(m+1)} &= \frac{\sum_{i=1}^{N} \tau(z_{ik}) \boldsymbol{X_i}}{\sum_{i=1}^{N} \tau(z_{ik})}, \\
\Sigma_k{}^{(m+1)} &= \frac{\sum_{i=1}^{N} \tau(z_{ik}) (\boldsymbol{X_i} - \boldsymbol{\mu}_k^{(m+1)})^T (\boldsymbol{X_i} - \boldsymbol{\mu}_k^{(m+1)})}{\sum_{i=1}^{N} \tau(z_{ik})}.
\end{aligned}$$

Repeat the above iterations until convergence.

#### 1.3 IMIX-Cor-Restrict

We conduct the EM algorithm similarly to IMIX-Cor with an restriction imposed on $\boldsymbol{\mu_k}$, we define

$$\boldsymbol{\Sigma_k^{-1}} = \begin{pmatrix} a_{11}^{(k)} & a_{12}^{(k)} & a_{13}^{(k)} \\ a_{12}^{(k)} & a_{22}^{(k)} & a_{23}^{(k)} \\ a_{13}^{(k)} & a_{23}^{(k)} & a_{33}^{(k)} \end{pmatrix}, \quad k = 1, \cdots, 8.$$

then $\boldsymbol{\mu}_k$ can be updated as:

$$
\begin{aligned}
\mu_{10}^{(m+1)} =& \frac{\sum_{i=1}^{N} X_{i1}\left(a_{11}^{(1)}\tau(z_{i1}) + a_{11}^{(3)}\tau(z_{i3}) + a_{11}^{(4)}\tau(z_{i4}) + a_{11}^{(7)}\tau(z_{i7})\right)}{\sum_{i=1}^{N}\left(a_{11}^{(1)}\tau(z_{i1}) + a_{11}^{(3)}\tau(z_{i3}) + a_{11}^{(4)}\tau(z_{i4}) + a_{11}^{(7)}\tau(z_{i7})\right)} \\
&+ \frac{\sum_{i=1}^{N}\left((X_{i2} - \mu_{20}^{(m)})(a_{12}^{(1)}\tau(z_{i1}) + a_{12}^{(4)}\tau(z_{i4})) + (X_{i2} - \mu_{21}^{(m)})(a_{12}^{(3)}\tau(z_{i3}) + a_{12}^{(7)}\tau(z_{i7}))\right)}{\sum_{i=1}^{N}\left(a_{11}^{(1)}\tau(z_{i1}) + a_{11}^{(3)}\tau(z_{i3}) + a_{11}^{(4)}\tau(z_{i4}) + a_{11}^{(7)}\tau(z_{i7})\right)} \\
&+ \frac{\sum_{i=1}^{N}\left((X_{i3} - \mu_{30}^{(m)})(a_{13}^{(1)}\tau(z_{i1}) + a_{13}^{(3)}\tau(z_{i3})) + (X_{i3} - \mu_{31}^{(m)})(a_{13}^{(4)}\tau(z_{i4}) + a_{13}^{(7)}\tau(z_{i7}))\right)}{\sum_{i=1}^{N}\left(a_{11}^{(1)}\tau(z_{i1}) + a_{11}^{(3)}\tau(z_{i3}) + a_{11}^{(4)}\tau(z_{i4}) + a_{11}^{(7)}\tau(z_{i7})\right)}, \\
\mu_{20}^{(m+1)} =& \frac{\sum_{i=1}^{N} X_{i2}\left(a_{22}^{(1)}\tau(z_{i1}) + a_{22}^{(2)}\tau(z_{i2}) + a_{22}^{(4)}\tau(z_{i4}) + a_{22}^{(6)}\tau(z_{i6})\right)}{\sum_{i=1}^{N}\left(a_{22}^{(1)}\tau(z_{i1}) + a_{22}^{(2)}\tau(z_{i2}) + a_{22}^{(4)}\tau(z_{i4}) + a_{22}^{(6)}\tau(z_{i6})\right)} \\
&+ \frac{\sum_{i=1}^{N}\left((X_{i1} - \mu_{10}^{(m)})(a_{12}^{(1)}\tau(z_{i1}) + a_{12}^{(4)}\tau(z_{i4})) + (X_{i1} - \mu_{11}^{(m)})(a_{12}^{(2)}\tau(z_{i2}) + a_{12}^{(6)}\tau(z_{i6}))\right)}{\sum_{i=1}^{N}\left(a_{22}^{(1)}\tau(z_{i1}) + a_{22}^{(2)}\tau(z_{i2}) + a_{22}^{(4)}\tau(z_{i4}) + a_{22}^{(6)}\tau(z_{i6})\right)} \\
&+ \frac{\sum_{i=1}^{N}\left((X_{i3} - \mu_{30}^{(m)})(a_{23}^{(1)}\tau(z_{i1}) + a_{23}^{(2)}\tau(z_{i2})) + (X_{i3} - \mu_{31}^{(m)})(a_{23}^{(4)}\tau(z_{i4}) + a_{23}^{(6)}\tau(z_{i6}))\right)}{\sum_{i=1}^{N}\left(a_{22}^{(1)}\tau(z_{i1}) + a_{22}^{(2)}\tau(z_{i2}) + a_{22}^{(4)}\tau(z_{i4}) + a_{22}^{(6)}\tau(z_{i6})\right)}, \\
\mu_{30}^{(m+1)} =& \frac{\sum_{i=1}^{N} X_{i3}\left(a_{33}^{(1)}\tau(z_{i1}) + a_{33}^{(2)}\tau(z_{i2}) + a_{33}^{(3)}\tau(z_{i3}) + a_{33}^{(5)}\tau(z_{i5})\right)}{\sum_{i=1}^{N}\left(a_{33}^{(1)}\tau(z_{i1}) + a_{33}^{(2)}\tau(z_{i2}) + a_{33}^{(3)}\tau(z_{i3}) + a_{33}^{(5)}\tau(z_{i5})\right)} \\
&+ \frac{\sum_{i=1}^{N}\left((X_{i1} - \mu_{10}^{(m)})(a_{13}^{(1)}\tau(z_{i1}) + a_{13}^{(3)}\tau(z_{i3})) + (X_{i1} - \mu_{11}^{(m)})(a_{13}^{(2)}\tau(z_{i2}) + a_{13}^{(5)}\tau(z_{i5}))\right)}{\sum_{i=1}^{N}\left(a_{33}^{(1)}\tau(z_{i1}) + a_{33}^{(2)}\tau(z_{i2}) + a_{33}^{(3)}\tau(z_{i3}) + a_{33}^{(5)}\tau(z_{i5})\right)} \\
&+ \frac{\sum_{i=1}^{N}\left((X_{i2} - \mu_{20}^{(m)})(a_{23}^{(1)}\tau(z_{i1}) + a_{23}^{(2)}\tau(z_{i2})) + (X_{i2} - \mu_{21}^{(m)})(a_{23}^{(3)}\tau(z_{i3}) + a_{23}^{(5)}\tau(z_{i5}))\right)}{\sum_{i=1}^{N}\left(a_{33}^{(1)}\tau(z_{i1}) + a_{33}^{(2)}\tau(z_{i2}) + a_{33}^{(3)}\tau(z_{i3}) + a_{33}^{(5)}\tau(z_{i5})\right)}, \\
\mu_{11}^{(m+1)} =& \frac{\sum_{i=1}^{N} X_{i1}\left(a_{11}^{(2)}\tau(z_{i2}) + a_{11}^{(5)}\tau(z_{i5}) + a_{11}^{(6)}\tau(z_{i6}) + a_{11}^{(8)}\tau(z_{i8})\right)}{\sum_{i=1}^{N}\left(a_{11}^{(2)}\tau(z_{i2}) + a_{11}^{(5)}\tau(z_{i5}) + a_{11}^{(6)}\tau(z_{i6}) + a_{11}^{(8)}\tau(z_{i8})\right)} \\
&+ \frac{\sum_{i=1}^{N}\left((X_{i2} - \mu_{20}^{(m)})(a_{12}^{(2)}\tau(z_{i2}) + a_{12}^{(6)}\tau(z_{i6})) + (X_{i2} - \mu_{21}^{(m)})(a_{12}^{(5)}\tau(z_{i5}) + a_{12}^{(8)}\tau(z_{i8}))\right)}{\sum_{i=1}^{N}\left(a_{11}^{(2)}\tau(z_{i2}) + a_{11}^{(5)}\tau(z_{i5}) + a_{11}^{(6)}\tau(z_{i6}) + a_{11}^{(8)}\tau(z_{i8})\right)} \\
&+ \frac{\sum_{i=1}^{N}\left((X_{i3} - \mu_{30}^{(m)})(a_{13}^{(2)}\tau(z_{i2}) + a_{13}^{(5)}\tau(z_{i5})) + (X_{i3} - \mu_{31}^{(m)})(a_{13}^{(6)}\tau(z_{i6}) + a_{13}^{(8)}\tau(z_{i8}))\right)}{\sum_{i=1}^{N}\left(a_{11}^{(2)}\tau(z_{i2}) + a_{11}^{(5)}\tau(z_{i5}) + a_{11}^{(6)}\tau(z_{i6}) + a_{11}^{(8)}\tau(z_{i8})\right)}, \\
\mu_{21}^{(m+1)} =& \frac{\sum_{i=1}^{N} X_{i2}\left(a_{22}^{(3)}\tau(z_{i3}) + a_{22}^{(5)}\tau(z_{i5}) + a_{22}^{(7)}\tau(z_{i7}) + a_{22}^{(8)}\tau(z_{i8})\right)}{\sum_{i=1}^{N}\left(a_{22}^{(3)}\tau(z_{i3}) + a_{22}^{(5)}\tau(z_{i5}) + a_{22}^{(7)}\tau(z_{i7}) + a_{22}^{(8)}\tau(z_{i8})\right)} \\
&+ \frac{\sum_{i=1}^{N}\left((X_{i1} - \mu_{10}^{(m)})(a_{12}^{(3)}\tau(z_{i3}) + a_{12}^{(7)}\tau(z_{i7})) + (X_{i1} - \mu_{11}^{(m)})(a_{12}^{(5)}\tau(z_{i5}) + a_{12}^{(8)}\tau(z_{i8}))\right)}{\sum_{i=1}^{N}\left(a_{22}^{(3)}\tau(z_{i3}) + a_{22}^{(5)}\tau(z_{i5}) + a_{22}^{(7)}\tau(z_{i7}) + a_{22}^{(8)}\tau(z_{i8})\right)} \\
&+ \frac{\sum_{i=1}^{N}\left((X_{i3} - \mu_{30}^{(m)})(a_{23}^{(3)}\tau(z_{i3}) + a_{23}^{(5)}\tau(z_{i5})) + (X_{i3} - \mu_{31}^{(m)})(a_{23}^{(7)}\tau(z_{i7}) + a_{23}^{(8)}\tau(z_{i8}))\right)}{\sum_{i=1}^{N}\left(a_{22}^{(3)}\tau(z_{i3}) + a_{22}^{(5)}\tau(z_{i5}) + a_{22}^{(7)}\tau(z_{i7}) + a_{22}^{(8)}\tau(z_{i8})\right)}, \\
\mu_{31}^{(m+1)} =& \frac{\sum_{i=1}^{N} X_{i3}\left(a_{33}^{(4)}\tau(z_{i4}) + a_{33}^{(6)}\tau(z_{i6}) + a_{33}^{(7)}\tau(z_{i7}) + a_{33}^{(8)}\tau(z_{i8})\right)}{\sum_{i=1}^{N}\left(a_{33}^{(4)}\tau(z_{i4}) + a_{33}^{(6)}\tau(z_{i6}) + a_{33}^{(7)}\tau(z_{i7}) + a_{33}^{(8)}\tau(z_{i8})\right)} \\
&+ \frac{\sum_{i=1}^{N}\left((X_{i1} - \mu_{10}^{(m)})(a_{13}^{(4)}\tau(z_{i4}) + a_{13}^{(7)}\tau(z_{i7})) + (X_{i1} - \mu_{11}^{(m)})(a_{13}^{(6)}\tau(z_{i6}) + a_{13}^{(8)}\tau(z_{i8}))\right)}{\sum_{i=1}^{N}\left(a_{33}^{(4)}\tau(z_{i4}) + a_{33}^{(6)}\tau(z_{i6}) + a_{33}^{(7)}\tau(z_{i7}) + a_{33}^{(8)}\tau(z_{i8})\right)} \\
&+ \frac{\sum_{i=1}^{N}\left((X_{i2} - \mu_{20}^{(m)})(a_{23}^{(4)}\tau(z_{i4}) + a_{23}^{(6)}\tau(z_{i6})) + (X_{i2} - \mu_{21}^{(m)})(a_{23}^{(7)}\tau(z_{i7}) + a_{23}^{(8)}\tau(z_{i8}))\right)}{\sum_{i=1}^{N}\left(a_{33}^{(4)}\tau(z_{i4}) + a_{33}^{(6)}\tau(z_{i6}) + a_{33}^{(7)}\tau(z_{i7}) + a_{33}^{(8)}\tau(z_{i8})\right)}.
\end{aligned}
$$

For two data types, we define

$$\boldsymbol{\Sigma_k^{-1}} = \begin{pmatrix} a_{11}^{(k)} & a_{12}^{(k)} \\ a_{12}^{(k)} & a_{22}^{(k)} \end{pmatrix}, \quad k = 1, \cdots, 4$$

then $\boldsymbol{\mu}_k$ can be updated as:

$$\begin{aligned}
\mu_{10}^{(m+1)} =& \frac{\sum_{i=1}^N X_{i1} \left( a_{11}^{(1)} \tau(z_{i1}) + a_{11}^{(3)} \tau(z_{i3}) \right)}{\sum_{i=1}^N \left( a_{11}^{(1)} \tau(z_{i1}) + a_{11}^{(3)} \tau(z_{i3}) \right)} \\
&+ \frac{\sum_{i=1}^N \left( (X_{i2} - \mu_{20}^{(m)}) a_{12}^{(1)} \tau(z_{i1}) + (X_{i2} - \mu_{21}^{(m)}) a_{12}^{(3)} \tau(z_{i3}) \right)}{\sum_{i=1}^N \left( a_{11}^{(1)} \tau(z_{i1}) + a_{11}^{(3)} \tau(z_{i3}) \right)} \\
\mu_{20}^{(m+1)} =& \frac{\sum_{i=1}^N X_{i2} \left( a_{22}^{(1)} \tau(z_{i1}) + a_{22}^{(2)} \tau(z_{i2}) \right)}{\sum_{i=1}^N \left( a_{22}^{(1)} \tau(z_{i1}) + a_{22}^{(2)} \tau(z_{i2}) \right)} \\
&+ \frac{\sum_{i=1}^N \left( (X_{i1} - \mu_{10}^{(m)}) a_{12}^{(1)} \tau(z_{i1}) + (X_{i1} - \mu_{11}^{(m)}) a_{12}^{(2)} \tau(z_{i2}) \right)}{\sum_{i=1}^N \left( a_{22}^{(1)} \tau(z_{i1}) + a_{22}^{(2)} \tau(z_{i2}) \right)} \\
\mu_{11}^{(m+1)} =& \frac{\sum_{i=1}^N X_{i1} \left( a_{11}^{(2)} \tau(z_{i2}) + a_{11}^{(4)} \tau(z_{i4}) \right)}{\sum_{i=1}^N \left( a_{11}^{(2)} \tau(z_{i2}) + a_{11}^{(4)} \tau(z_{i4}) \right)} \\
&+ \frac{\sum_{i=1}^N \left( (X_{i2} - \mu_{20}^{(m)}) a_{12}^{(2)} \tau(z_{i2}) + (X_{i2} - \mu_{21}^{(m)}) a_{12}^{(4)} \tau(z_{i4}) \right)}{\sum_{i=1}^N \left( a_{11}^{(2)} \tau(z_{i2}) + a_{11}^{(4)} \tau(z_{i4}) \right)} \\
\mu_{21}^{(m+1)} =& \frac{\sum_{i=1}^N X_{i2} \left( a_{22}^{(3)} \tau(z_{i3}) + a_{22}^{(4)} \tau(z_{i4}) \right)}{\sum_{i=1}^N \left( a_{22}^{(3)} \tau(z_{i3}) + a_{22}^{(4)} \tau(z_{i4}) \right)} \\
&+ \frac{\sum_{i=1}^N \left( (X_{i1} - \mu_{10}^{(m)}) a_{12}^{(3)} \tau(z_{i3}) + (X_{i1} - \mu_{11}^{(m)}) a_{12}^{(4)} \tau(z_{i4}) \right)}{\sum_{i=1}^N \left( a_{22}^{(3)} \tau(z_{i3}) + a_{22}^{(4)} \tau(z_{i4}) \right)}.
\end{aligned}$$

### 2 Simulation Studies

#### 2.1 Simulation Parameters Mimicking TCGA Bladder Cancer Real Data

The parameters used for multivariate normal mixture for simulation studies scenario 6 mimicking TCGA bladder cancer real data are shown here, $\hat{\pi}$=(0.268577053, 0.233897556, 0.189871018, 0.006866345, 0.256186213, 0.008889446, 0.010178071, 0.025534298), the mean vectors for the eight components are

$\boldsymbol{\mu_1} = (0.5012989, 0.3832367, 0.4550859)$; $\boldsymbol{\mu_2} = (3.6045370, 0.3832367, 0.4550859)$;
$\boldsymbol{\mu_3} = (0.5012989, 4.2595606, 0.4550859)$; $\boldsymbol{\mu_4} = (0.5012989, 0.3832367, 4.2136909)$;
$\boldsymbol{\mu_5} = (3.6045370, 4.2595606, 0.4550859)$; $\boldsymbol{\mu_6} = (3.6045370, 0.3832367, 4.2136909)$;
$\boldsymbol{\mu_7} = (0.5012989, 4.2595606, 4.2136909)$; $\boldsymbol{\mu_8} = (3.604537, 4.259561, 4.213691)$.

The covariance matrices for the eight components are

$$\boldsymbol{\Sigma_1} = \begin{pmatrix} 1.26 & 0.01 & 0.05 \\ 0.01 & 1.04 & 0.02 \\ 0.05 & 0.02 & 1.21 \end{pmatrix}; \boldsymbol{\Sigma_2} = \begin{pmatrix} 1.50 & -0.01 & -0.02 \\ -0.01 & 1.02 & -0.04 \\ -0.02 & -0.04 & 1.14 \end{pmatrix}; \boldsymbol{\Sigma_3} = \begin{pmatrix} 1.21 & 0.09 & 0.02 \\ 0.09 & 2.52 & -0.02 \\ 0.02 & -0.02 & 1.22 \end{pmatrix};$$

$$\boldsymbol{\Sigma_4} = \begin{pmatrix} 0.90 & 0.01 & 0.06 \\ 0.01 & 1.01 & 0.05 \\ 0.06 & 0.05 & 1.12 \end{pmatrix}; \boldsymbol{\Sigma_5} = \begin{pmatrix} 1.84 & 0.22 & -0.09 \\ 0.22 & 3.58 & -0.01 \\ -0.09 & -0.01 & 1.21 \end{pmatrix}; \boldsymbol{\Sigma_6} = \begin{pmatrix} 1.92 & -0.04 & 0.16 \\ -0.04 & 1.05 & -0.16 \\ 0.16 & -0.16 & 1.23 \end{pmatrix};$$

$$\mathbf{\Sigma_7} = \begin{pmatrix} 1.01 & 0.11 & 0.13 \\ 0.11 & 2.80 & 0.36 \\ 0.13 & 0.36 & 1.24 \end{pmatrix} ; \mathbf{\Sigma_8} = \begin{pmatrix} 1.63 & 0.29 & 0.08 \\ 0.29 & 4.02 & 0.09 \\ 0.08 & 0.09 & 1.39 \end{pmatrix} .$$

#### 2.2 Model Calibration Estimation

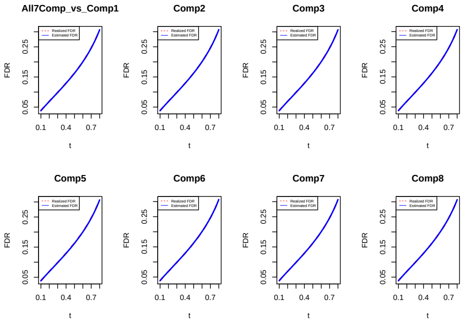


(a) Scenario1

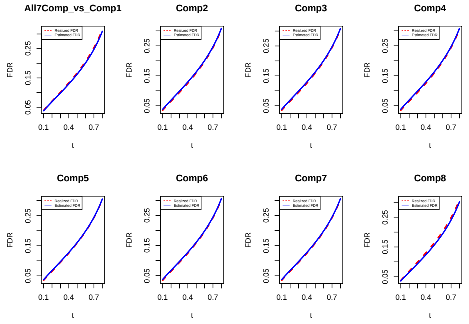


(b) Scenario2

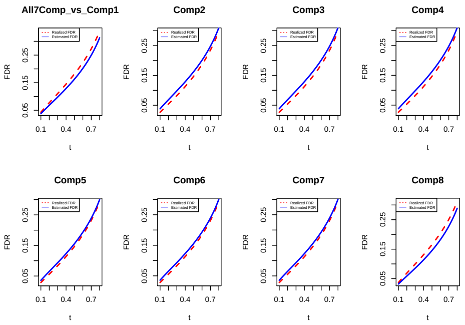


(c) Scenario3

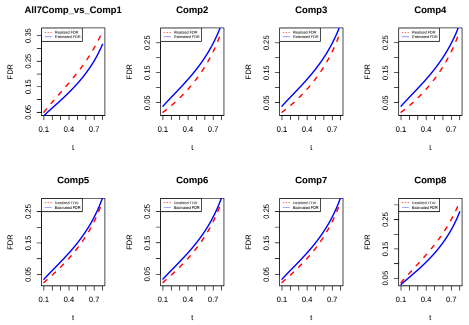


(d) Scenario4

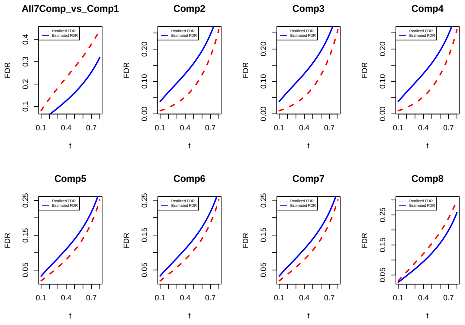


(e) Scenario5

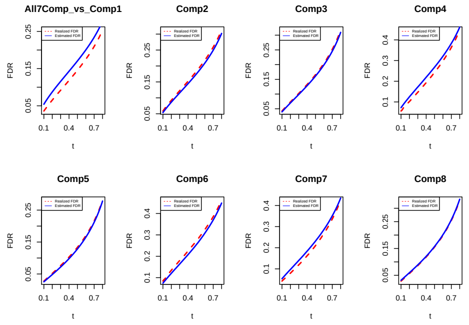


(f) Scenario6

Figure 1: Model calibration of IMIX-Ind for 1000 simulation results.

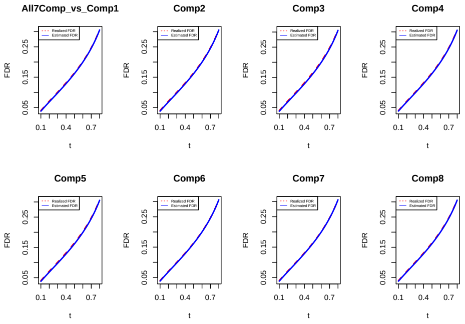


(a) Scenario1

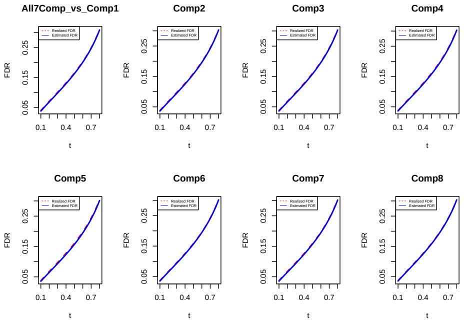


(b) Scenario2

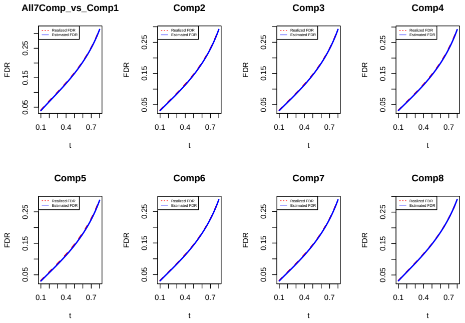


(c) Scenario3

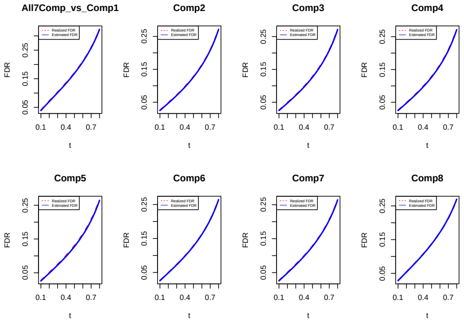


(d) Scenario4

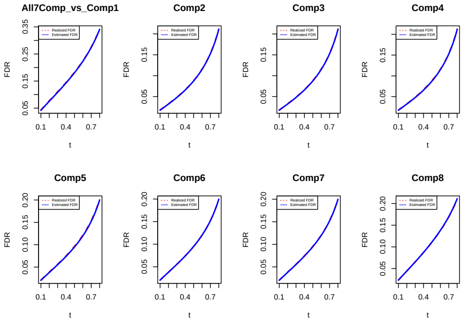


(e) Scenario5

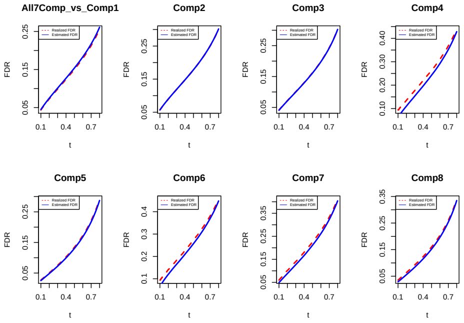


(f) Scenario6

Figure 2: Model calibration of IMIX-Cor for 1000 simulation results.

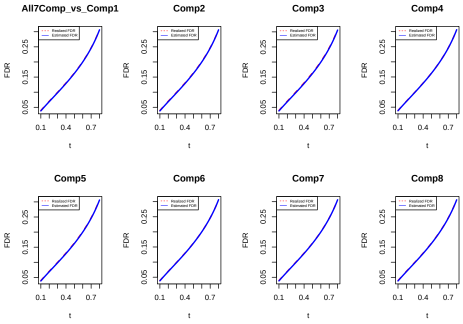


(a) Scenario1

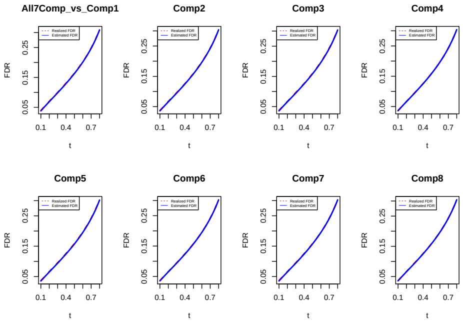


(b) Scenario2

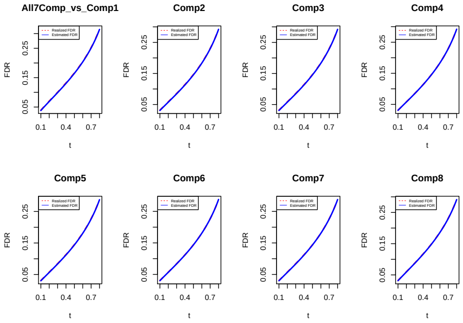


(c) Scenario3

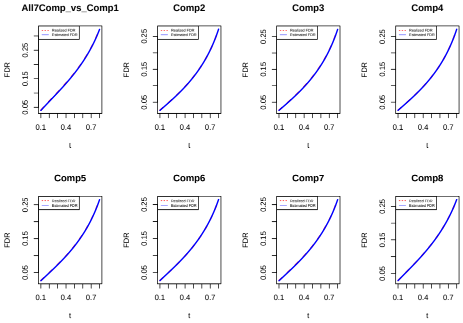


(d) Scenario4

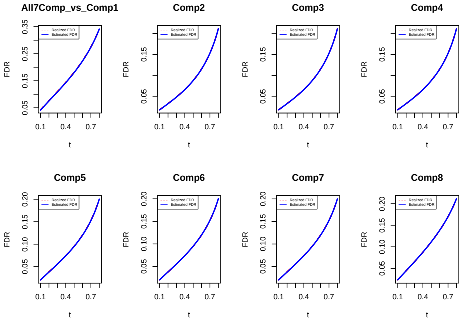


(e) Scenario5

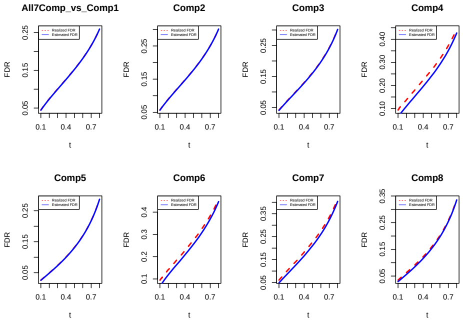


(f) Scenario6

Figure 3: Model calibration of IMIX-Cor-Restrict for 1000 simulation results.

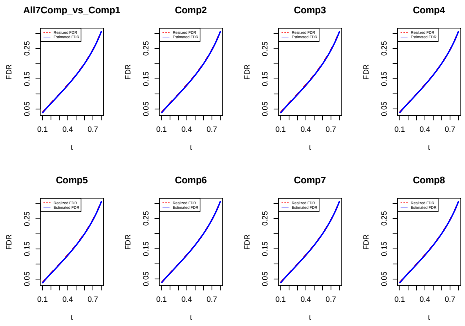


(a) Scenario1

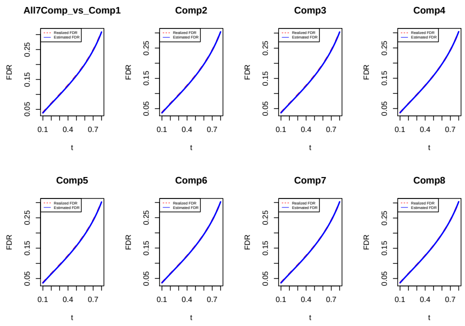


(b) Scenario2

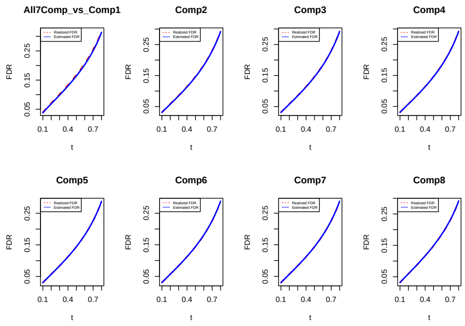


(c) Scenario3

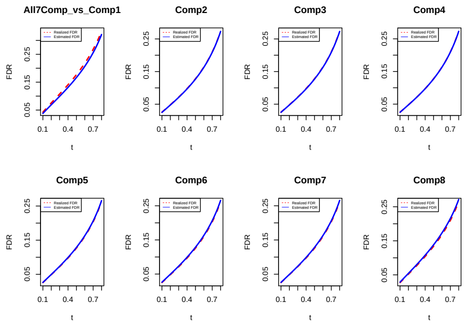


(d) Scenario4

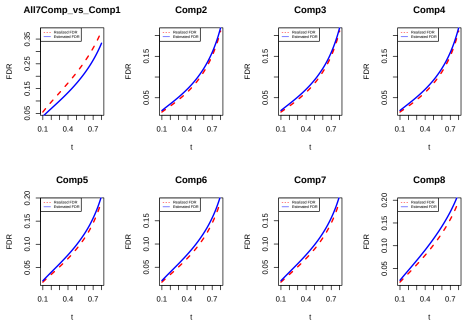


(e) Scenario5

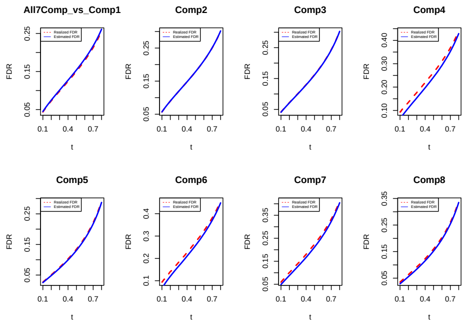


(f) Scenario6

Figure 4: Model calibration of IMIX-Cor-Twostep for 1000 simulation results.

#### 2.3 Model Selection

| Number of Components | Balanced | Unbalanced |
|---|---|---|
| 7 Components | (0.25,0.125,0.125,0.125,0.125,0.125,0.125) | (0.304,0.095,0.290,0.017,0.269,0.004,0.021) |
| 8 Components | (0.125,0.125,0.125,0.125,0.125,0.125,0.125,0.125) | (0.300,0.095,0.290,0.017,0.269,0.004,0.021,0.004) |

Table 1: Mixing proportions in simulation study setup 2.

Let $x_i$ be the group label, here we let it be from 1 to 8, $x_i = 1, \cdots, 8$. a is the slope, b is the intercept. Let

$$a \sum_{i=1}^{8} x_i + b = 1$$

Now we consider to solve this equation

$$\begin{cases} 36a + 8b = 1 \\ 8a + b = c \end{cases} \tag{1}$$

Here, we let $c = 0.005, 0.01, 0.05, 0.1$. Solve the equation and get the proportions for each component for the 4 scenarios, the proportion of component $i$ would be $ax_i + b$.

| | Proportion |
|---|---|
| c=0.1 | (0.150,0.143,0.136,0.129,0.121,0.114,0.107,0.100) |
| c=0.05 | (0.200,0.179,0.157,0.136,0.114,0.093,0.071,0.05) |
| c=0.01 | (0.240,0.207,0.174,0.141,0.109,0.076,0.043,0.010) |
| c=0.005 | (0.245,0.211,0.176,0.142,0.108,0.074,0.039,0.005) |

Table 2: Mixing proportions in simulation study setup 3.

| | Proportion |
|---|---|
| c=0.1 | (0.129,0.129,0.129,0.129,0.128,0.128,0.128,0.100) |
| c=0.05 | (0.136,0.136,0.136,0.136,0.136,0.135,0.135,0.05) |
| c=0.01 | (0.144,0.141,0.141,0.141,0.141,0.141,0.141,0.010) |
| c=0.005 | (0.143,0.142,0.142,0.142,0.142,0.142,0.142,0.005) |

Table 3: Mixing proportions in simulation study setup 4, one group unbalance.

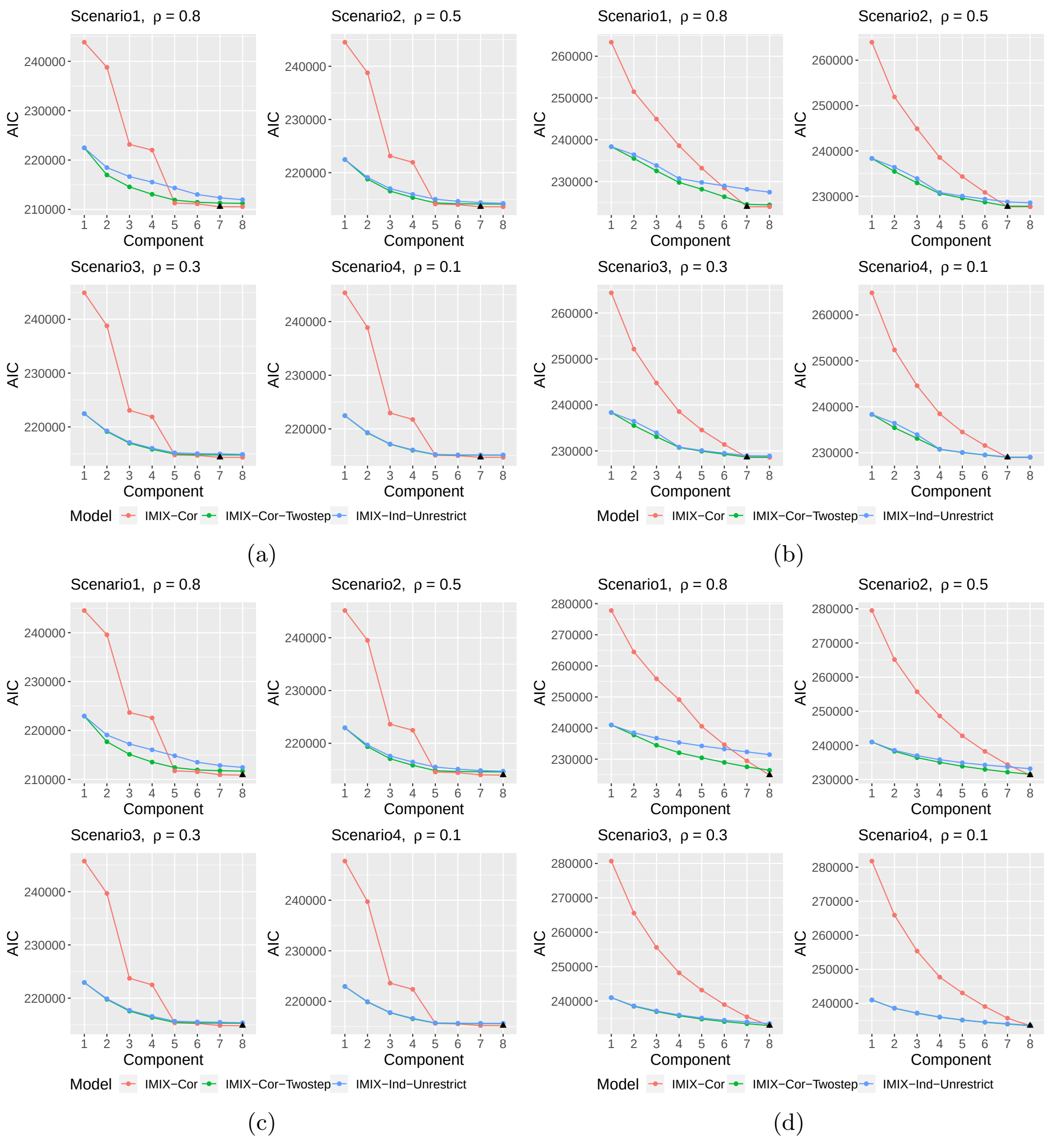


Figure 5: Simulation study on selecting the number of mixture components using AIC. (a)Unbalanced setting, 7-component mixture model; (b)Balanced setting, 7-component mixture model; (c)Unbalanced setting, 8-component mixture model; (d)Balanced setting, 8-component mixture model. Black triangle represents the model AIC selects. $\rho$ is the correlation between data types.

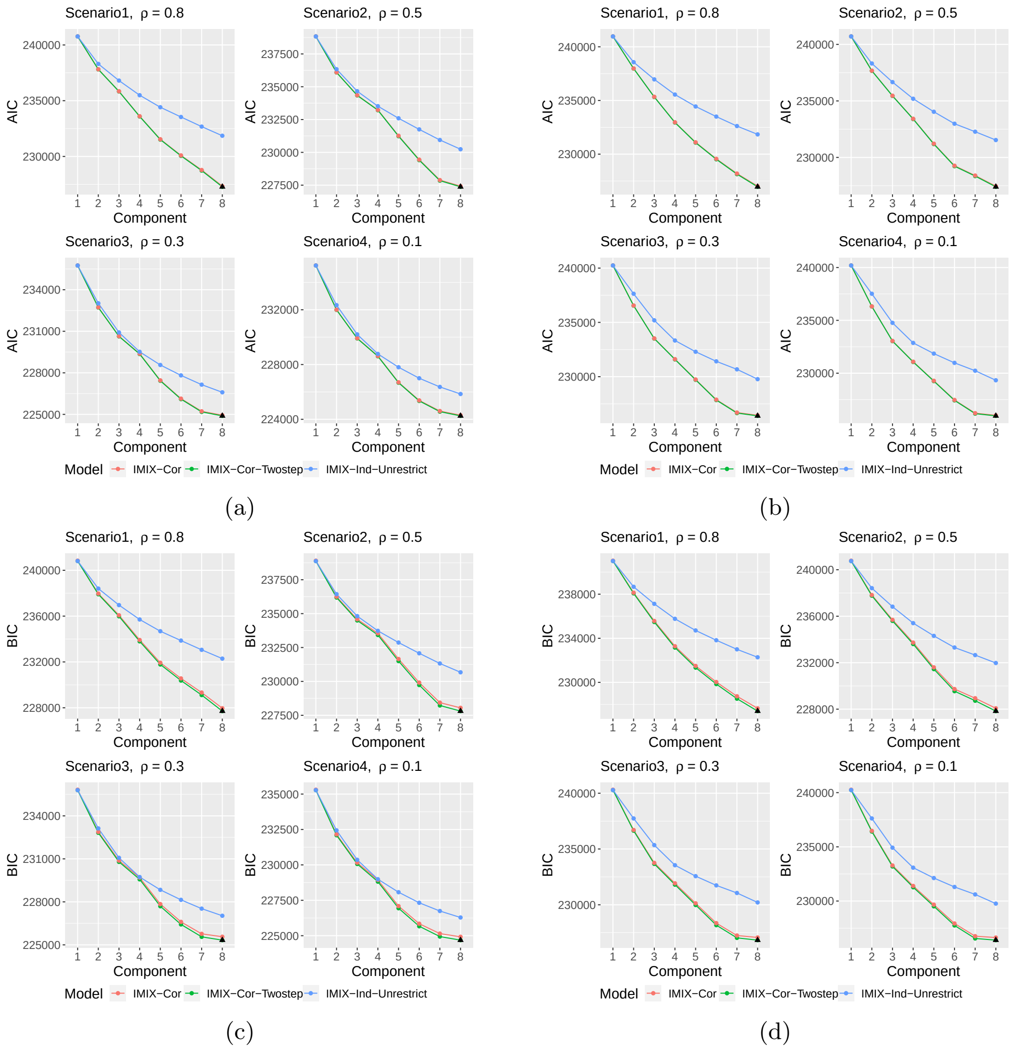


Figure 6: Model selection of AIC and BIC when the mixing proportions are unbalanced. AIC: (a) simulation set up 3; (b) simulation set up 4; BIC: (c) simulation set up 3; (d) simulation set up 4. Black triangle represents the model AIC/BIC selects. $\rho$ is the correlation between data types.

##### 2.4 Computational Time of IMIX

| Computation time (in seconds) | Mean | SD |
|---|---|---|
| IMIX-Ind | 4.501 | 0.879 |
| IMIX-Cor | 970.901 | 167.922 |
| IMIX-Cor-Restrict | 417.531 | 57.425 |
| IMIX-Cor-Twostep | 217.379 | 25.787 |

Table 4: Computational time needed (mean and standard deviation (SD) with 20 000 genes of $H = 3$ data types) for IMIX models under comparison in the simulation study 1 (Figure 1; Scenario 3 with inter-data correlation of 0.3).

| Number of iteration | Mean | SD |
|---|---|---|
| IMIX-Ind | 67 | 8.652 |
| IMIX-Cor | 161 | 24.606 |
| IMIX-Cor-Restrict | 71 | 7.597 |
| IMIX-Cor-Twostep | 42 | 3.497 |

Table 5: Number of iterations needed (mean and standard deviation (SD) with 20 000 genes of $H = 3$ data types) for IMIX models under comparison in the simulation study 1 (Figure 1; Scenario 3 with inter-data correlation of 0.3).

#### 3 Real Data Applications to The Cancer Genome Atlas (TCGA)

##### 3.1 Data Preprocessing and Quality Control

For bladder cancer in the TCGA, copy number variation (CNV) and methylation data were retrieved from TCGA2STAT (Wan et al., 2016), RNAseq data was retrieved from Broad Institute Genome Data Analysis Centers TCGA (http://gdac.broadinstitute.org/), and preprocessed and log transformed previously as described in Guo et al. (2019). All three datasets were with reference genome build hg19. CNV was array-based level-3 gene level data. Methylation data was measured on the Illumina Infinium HumanMethylation450 (450K)

BeadChip array at over 480,000 sites. The CpG sites with missing values in more than 10% of the total samples were filtered out. We excluded all the probes on the sex chromosomes, the probes of target polymorphic CpGs that overlaps with known SNPs (Fortin et al., 2017), and the probes that are cross-reactive (Chen et al., 2013). The methylation data was normalized using the Beta-Mixture Quantile (BMIQ) Normalization function from R package "wateRmelon" (Ruth, 2013). For each data type, we only included Caucassian origin samples and excluded samples with missing phenotypes. In the end, for downstream individual level analysis, there were 373 DNA methylation samples, 391 RNA-Seq samples, and 387 CNV samples with $N = 15\,672$ genes.

Individual level test was conducted with respect to the binary molecular subtypes using logistic regression adjusting for the clinical covariates, including age, sex, race, smoking status and pathologic stage. The same procedure was conducted for RNAseq and CNV data. For probe level methylation data, we conducted the set-based test within each gene, the sequence kernel association test (SKAT) (Wu et al., 2011). The summary statistics p value was collected for the three data types, and then transformed to Z scores for IMIX analysis.

For pancreatic cancer in the TCGA, we preprocessed the CNV data with the same quality control procedures as bladder cancer dataset. The level-3 RSEM RNAseq data was retrieved from TCGA2STAT (Wan et al., 2016) and log transformed. Same preprocessing procedure on the samples were performed. Individual level test was conducted for the time-to-event outcome using the Cox proportional hazards model to each of the 15 472 genes respectively on 157 RNA-Seq samples and 161 CNV samples adjusting for age, gender, and smoking status. The summary statistics p value was collected for the two data types, and then transformed to Z scores for IMIX analysis.

The molecular subtypes of bladder cancer patients in the TCGA were retrieved from previous work (Guo et al., 2019).

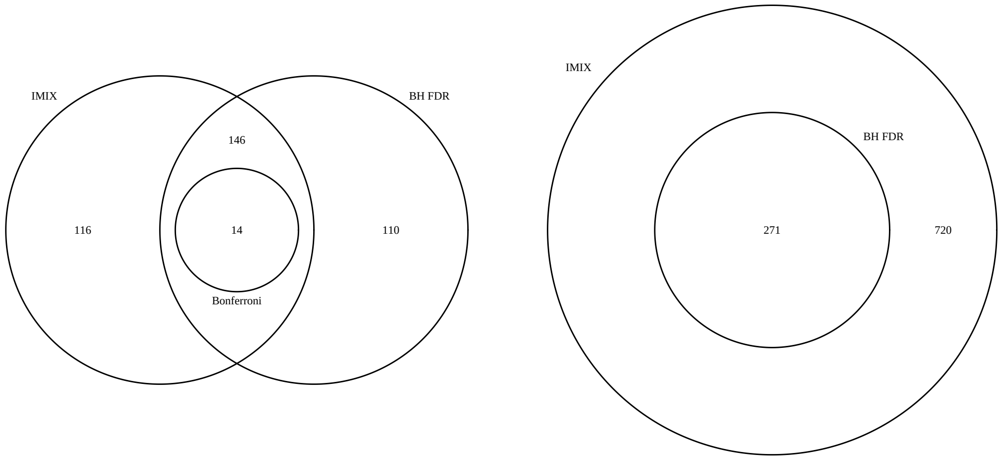


(a) (b)

Figure 7: Comparisons between the number of significant genes in the TCGA datasets detected by the IMIX framework, Benjamini-Hochberg FDR (BH-FDR), and Bonferroni correction. (a) Number of genes detected by the IMIX framework, BH-FDR, and Bonferroni correction that are associated with the molecular subtypes of muscle-invasive bladder cancer through DNA methylation, gene expression, and CNV, with adaptive FDR control at $\alpha = 0.2$, estimated $\widehat{\mathrm{mFDR}}_8 = 0.1995$. (b) Number of genes detected by IMIX and BH-FDR that are associated with the survival status of pancreatic cancer patients through gene expression and CNV, with adaptive FDR control at $\alpha = 0.2$, estimated $\widehat{\mathrm{mFDR}}_4 = 0.2$.

#### 3.2 The Directed Acyclic Graphs (DAGs) of 61 Significant Genes in Component 8

We estimated the causal relationships between DNA methylation, gene expression, and CNV of the 61 genes in component 8 by applying Bayesian networks (Scutari, 2017) with the target nominal type I error rate at 0.01. The directed acyclic graphs (DAGs) based on conditional independence tests with a restriction of causal direction from CNV to E showed six different patterns of causal structures for our data. In particular, seven genes had a full model with connections of CNV→E, E−M, M−CNV (Fig S8(1)); five genes showed causal effect of both CNV and DNA methylation on gene expression, while CNV and DNA methylation were marginally independent (Fig S8(2)); 17 genes had a reactive model as CNV→E→M (Fig S8(3)), which had also been reported by previous research (Sun et al., 2018). Seventeen genes showed that DNA methylation and gene expression were conditionally independent given CNV as M−CNV→E (Fig S8(4)) with the causal direction between CNV and DNA methylation indistinct from the statistical test. One gene showed conditional independence between CNV and gene expression given DNA methylation (Fig S8(5)); however, the direction of the DAG was not available due to Markov equavalence, i.e., all three scenarios ( CNV → M→E; E→M→CNV; CNV←M→E) led to conditional independence. This model was likely to be a true causal model as CNV→M→E. Four genes showed a dependence structure between gene expression and CNV given DNA methylation (Fig S8(6)). The rest of the 10 genes showed no triangular association pattern.

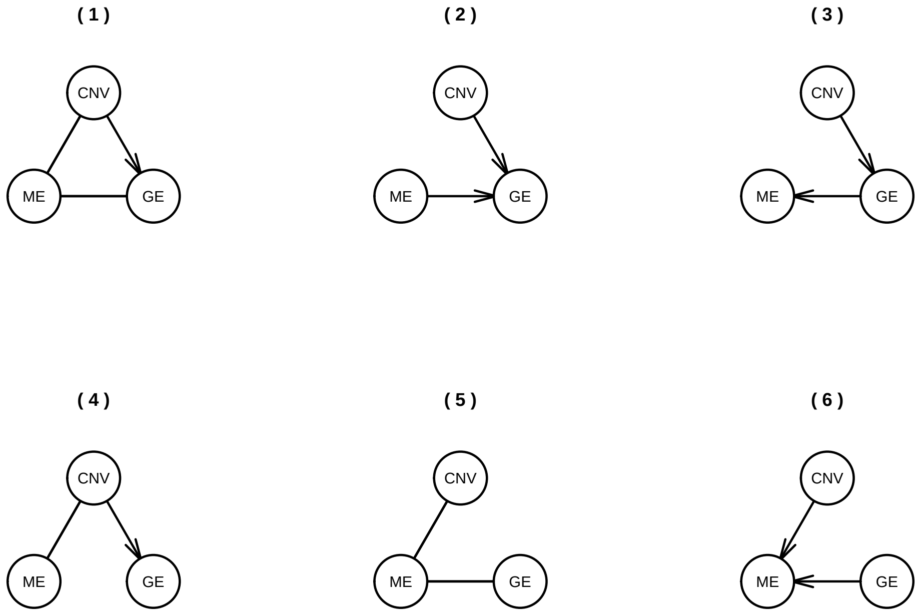


Figure 8: The Directed acyclic graphs (DAGs) based on Bayesian network with a restriction of causal direction from CNV to gene expression for the genes in component 8 of TCGA bladder cancer with across-data-type FDR controlled at $\alpha = 0.01$.

Analysis: bladder_fdr001_adaptive - 2020-02-20 12:21 PM

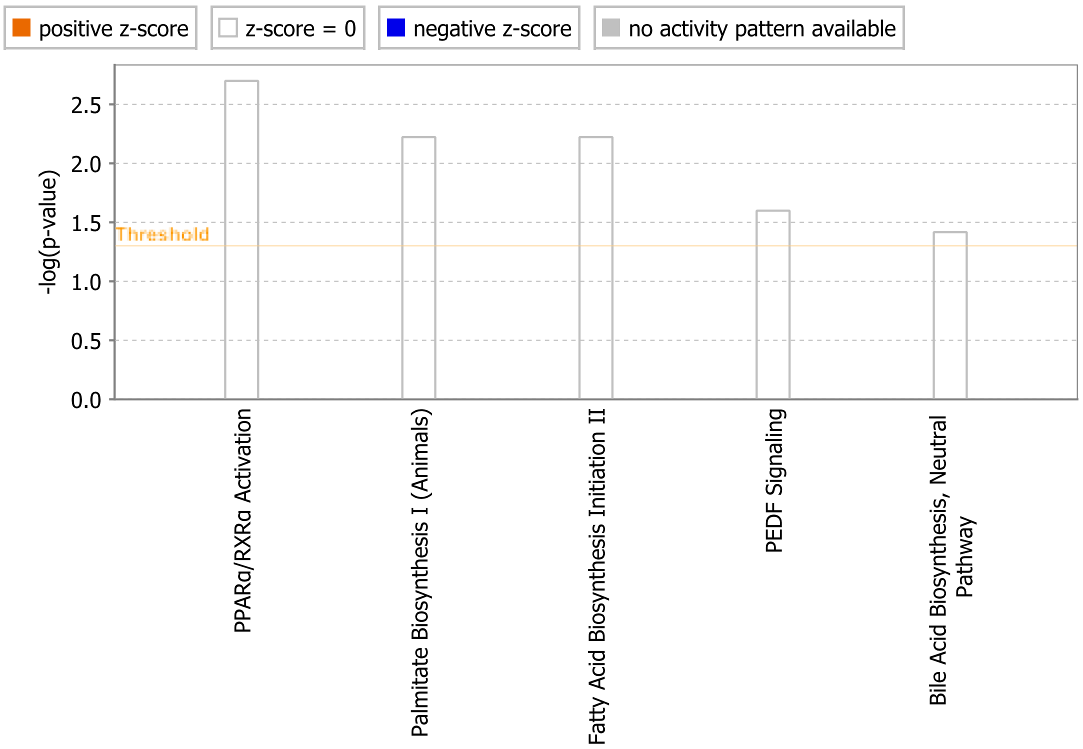


© 2000-2020 QIAGEN. All rights reserved.

Figure 9: The canonical pathways identified by the Ingenuity Pathway Analysis (IPA) on the 61 significant genes in component 8 with adaptive FDR controlled at $\alpha = 0.01$ of bladder cancer subtypes in the TCGA.
